# Wireless Electrical-Molecular Quantum Signalling for Cancer Cell Induced Death

**DOI:** 10.1101/2023.03.02.529075

**Authors:** Akhil Jain, Jonathan Gosling, Shaochuang Liu, Haowei Wang, Eloise M. Stone, Lluïsa Pérez-García, David B. Amabilino, Mark Fromhold, Stuart Smith, Ruman Rahman, Yitao Long, Lyudmila Turyanska, Frankie J. Rawson

## Abstract

Quantum biological tunnelling for electron transfer (QBET) is involved in controlling cellular behaviour. Control of electrical-molecular communication could revolutionise the development of disruptive technologies for understanding and modulating electrically induced molecular signalling. Current communication technology is not appropriate for interfacing with cells at a spatial/temporal level equivalent to the native biological signalling. This limits our ability to tune cell function by controlling single molecular events. Here, we merge wireless nano-electrochemical tools with cancer cells. Gold-bipolar nanoelectrodes functionalised with redox active species were developed as electric field stimulated bio-actuators, that we term bio-nanoantennae. We show that a remote electrical input regulates electron transport between the redox molecules on the bio-nanoantennae in a selective manner. The wireless modulation of electron transport results in QBET triggering apoptosis in patient-derived cancer cells, representing electrical-induced induced controlled molecular signalling. Transcriptomics data highlight the electric field-induced nanoantenna targets the cancer cells in a unique manner. The insight concerning action and functional nanomaterials opens a plethora of applications in healthcare. This approach may lead to new quantum-based medical diagnostics and treatments, as well as a fundamental understanding of biological physics.

---

Bioelectrical endogenous currents are essential for life. We are entering a new era where bioelectricity, defined as the electrical language of cells, is now realised to programme cell function.^1,2^ The cell is increasingly viewed as a mass of bioelectrical interconnected circuits.^3^ One of the most well-known electron transfer pathways is photosynthesis and was one of the first whose mechanism was linked to quantum mechanical effects.^4,5^ Quantum biology is still in its infancy, but future developments in biology and medicine are expected to be underpinned by an understanding of quantum mechanical processes.^6^ Cytochrome *c* (C) is an important redox-active protein that can induce apoptosis when its redox state is modulated through an electron transfer process at the heme.^7^ This process can occur via electron tunnelling.^8,9,10^ However, our ability to communicate electrically with such systems is limited by a mismatch in communication technology and selective targeting bio-interfacing of electrical actuation systems.

Remotely induced electrical-molecular communication (electrical input causing a molecularly targeted redox change which induces an alteration in cell behaviour) inside cells opens the possibility of creating new disruptive technologies, including the development of new quantum medicines for cancer treatments.^11^ However, on-demand targeted wireless electrical-molecular communication within cells is yet to be realised, mostly because of a lack of suitable communication technologies for interfacing with cells in a medium at a spatial/temporal level observed in native biological communication that occurs. Moreover, technological innovation in electrical transduction via bio-nanoantenna (defined as receiving an electrical input and converting to biological output) capable of receiving a remote externally applied electrical field input and converting this to bio-signalling events has not been achieved. The aim of this study was therefore to pioneer solutions to these challenges.

Gold nanoparticles^12^ and carbon nanotubes can behave as bipolar nanoelectrodes^13^ within cells when external electric fields (EFs) are applied. The bipolar nanoelectrodes become polarised when an EF is applied, leading to a voltage gradient across the particle. With a sufficiently large potential difference between the poles of the electrode, the thermodynamic driving force can cause electrochemically induced redox reactions to occur.^14,15^ It was thought that nanoscale wireless electrochemistry (also known as bipolar electrochemistry) was not possible in the presence of cells as high voltages were used.^16,17^ Importantly, most recently, bipolar electrodes in series led to a dramatic drop in cell impedance,^18^ which may allow electrical inputs to be used without inducing cell damage. Redox reactions at carbon nanotube porins, acting as bipolar nanoelectrodes, can occur at unprecedentedly low applied voltages in cells without inducing direct cell death.^19^ We, therefore, designed an approach that uses bi-functionalised bipolar nanoelectrodes with attached redox active molecules, we term bio-nanoantennae, in combination with applied EFs could be used to modulate electron transfer. This electronic signal is then converted into a molecular actuation in a specific targeted metabolic pathway (**Fig. 1**).

**Figure 1.**
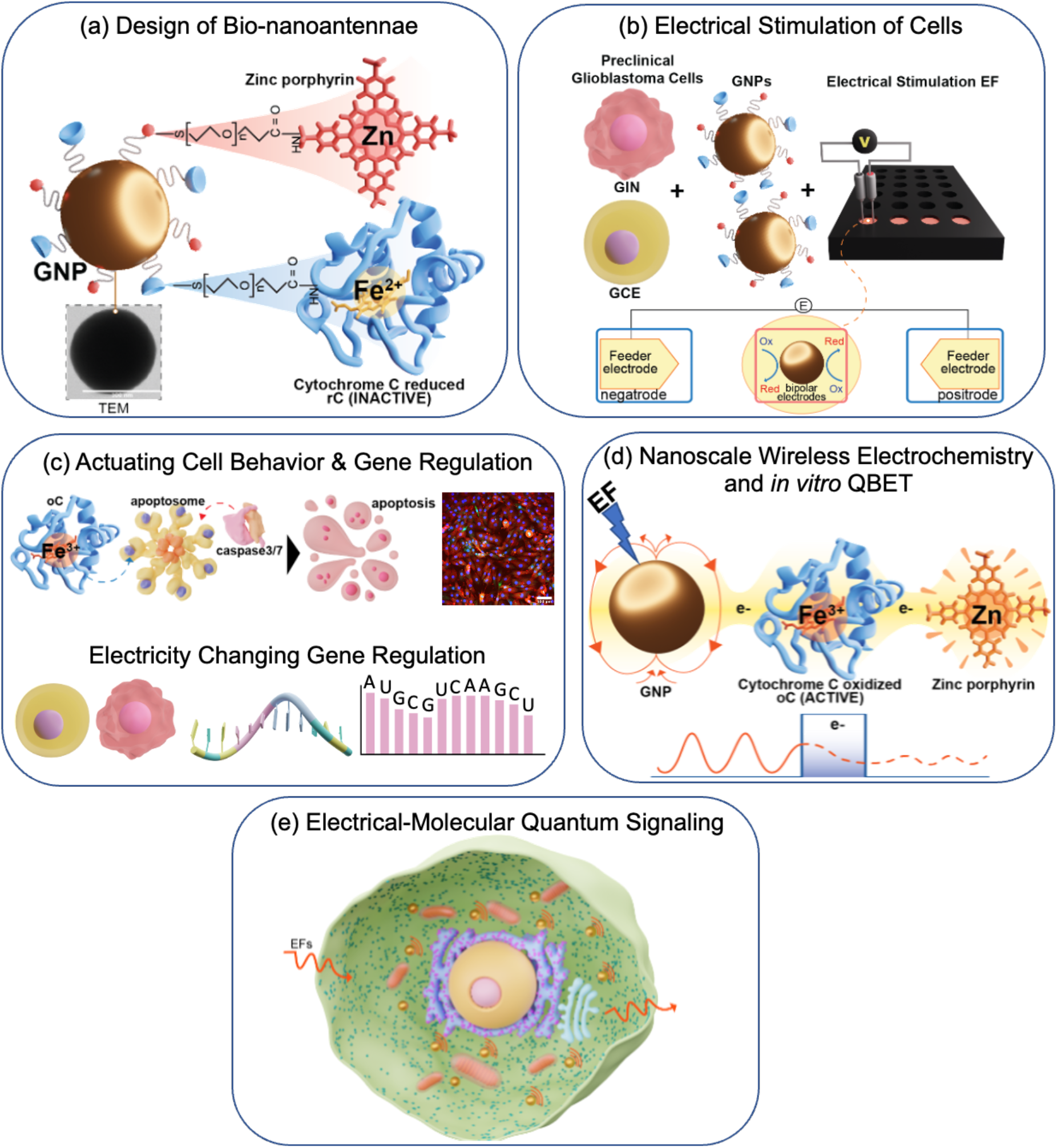
Illustration of AC-EF responsive bio-nanoantennae mediated wireless electrical-molecular quantum signalling to induce cell death. **(a)** Bio-nanoantennae were synthesised by covalently conjugating reduced C (rC) and Z to carboxylic PEG functionalised 100 nm citrate capped gold nanoparticles (GNP100) using EDC/NHS chemistry. **(b)** Primary patient-derived GBM cells viz. GIN (derived from the GBM infiltrative margin) and GCE (derived from the GBM proliferative core) cells were incubated with bio-nanoantennae (GNP100@rC@Z) to enable their uptake. These GBM cells were electrically stimulated (ES) with AC EFs of 3 MHz at 0.65 V/cm. **(c)** These applied AC EF caused the intracellular wireless electrochemistry to occur at bio-nanoantennae surface thus inducing caspase 3/7 mediated apoptosis of GBM cells. We examined the electrical-molecular signalling by connecting gene regulation with AC EFs mediated cell death in GBM cells. **(d)** AC EFs were applied to induce wireless electrochemistry at the surface of bio-nanoantennae to wirelessly switch the redox state of C (reduced to oxidised) and Z, ex-situ. Mathematical modelling and plasmon resonance scattering spectroscopy (PRS) were conducted to confirm QBET in the proposed system. **(e)** Diagrammatic representation to demonstrate AC EFs responsive bio-nanoantennae for Electrical-Molecular Quantum Signalling with GBM cells.

The strategy to fulfil the development of the first quantum electrical-molecular communication tool was multi-pronged. Firstly, we were inspired by the observation that C electron transfer is mediated through quantum biological electron tunnelling (QBET).^10^ Therefore, we functionalised gold nanoparticles (GNPs) with an electron donor C and a redox mediator in the form of zinc porphyrin (Z) and subject to electrical induction, therefore, resulting in an electronic bio-nanoantenna (Fig **1a**). We show that the electrical input confers specific signalling to cells via the bio-nanoantennae to induce Glioblastoma (GBM) cell death through apoptosis. The transcriptomic analysis enabled elucidation of the biochemical signalling when using the electrical-molecular communication tool (Fig **1b & 1c**). We show that on the application of an external electric field, wireless electrochemistry is induced at the nanoparticle surface switching the redox state of C. Next, we used multiple analytical approaches to show that the redox state of the C changes with a resonant electrical input of alternating current electric field (AC-EF) at 3 MHz using an applied voltage of 0.65V/cm. We propose that the QBET phenomenon and resonant electron transfer between the C and Z facilitate cellular apoptosis (Fig **1d**). The data suggest that electrical field molecular actuation in the treatment of a disease may represent a first-in-class quantum functional medicine tool (Fig **1e**). This could open exciting opportunities for the development of quantum nanomedicines and cancer treatments and provide a new means of modulating cell metabolism.

## Design of bio-nanoantennae for association and actuating patient derived GBM cell death

To electrically communicate with cells at an equivalent level to that underpinning molecular biochemistry, we initially used 100 nm PEGylated spherical gold nanoparticles (GNP100) as bipolar nanoelectrodes that can be functionalised to yield nanoantennae. These GNPs can sense electric fields applied extracellularly and drive surface redox reactions,^12,18^ which could modulate the redox state of surface-bound Z and C that would actuate a cell-specific signalling pathway, apoptosis. Apoptosis is mediated (in part) by C bioelectrochemistry in which the oxidised state (Fe^3+^) facilitates apoptosis-mediated cell death.^20^ Therefore, we functionalised GNP100 with an electrical donor i.e. reduced C (rC, inactive: Fe^2+^) and a redox mediator Z through carbodiimide coupling chemistry to form bio-nanoantennae (GNP100@rC@Z) (Figure **1a**). On application of an electric field, the particles would polarise providing the thermodynamic driving force to oxidize the rC.^21^ To validate our hypothesis and QBET, the effect of the NP size and the PEG linker length, 20 and 50 nm particles were also functionalized with rC and Z using PEG linker of varying molecular weights (1 KDa, 2 KDa, 3.5KDa, 5 kDa) to yield different size bio-nanoantennae (GNP20@rC@Z and GNP50@rC@Z).

The successful bifunctionalisation of PEGylated (2 kDa) 100 nm citrate capped GNPs (GNP100) with rC and Z is evident from the characterization of GNP100@rC@Z. The diameter of GNP100@rC@Z as analysed by TEM was found to be 105 ± 2 nm (Fig. **2a**). Dynamic Light Scattering (DLS) analysis indicated an average hydrodynamic diameter (*h*_d_) increase from 104.9 nm (PEGylated GNPs) to 118.8 nm for the GNP100@rC@Z, suggesting successful conjugation of rC and Z (Supplementary Fig. **1**). A change in zeta potential (ζ) of GNP100@rC@Z (-28.5 mV) compared to GNP100 (-27.7 mV), GNP100@rC (-14.8 mV), and GNP100@Z (-40 mV) suggested the bi-functionalisation process was successful (Supplementary Fig. **2**). UV-Vis absorption spectrum of GNP100@rC@Z before ES in PBS revealed a broad peak centred at 418 nm attributed to overlapping absorption of rC and Z (Fig. 2 b & c). This peak was deconvoluted and Marquardt fitting algorithm was applied,^23^ which revealed two components centred at 412 nm and 423 nm attributed to rC and Z (Supplementary Fig. **3a-b**), respectively, which were corroborated using UV-Vis spectrum of native rC and Z in PBS (Supplementary Fig. **3c**). The obtained peaks were used for the quantification of rC, and Z attached to each GNP100@rC@Z, which revealed homogenous and monolayer conjugation of rC and Z on GNP100 (Supplementary Fig. **3d-e** and Table **1-4**). Cyclic voltammetry was carried out to study the redox behaviour of rC and Z on a bifunctionalised system (Fig. **2d**). Two redox couples were observed in GNP100@rC@Z, which are attributed to C and Z, while on the other hand, the control samples only showed a redox couple corresponding to either C or Z, which have been characterised by others previously (Supplementary Fig. **4a-c**).^24,25^ The heterogeneous electron transfer rate coefficient (*k*0) of C for GNP100@rC was calculated to be 9.6 × 10^-^3 cm/s (Supplementary Table **5**), while that for GNP100@rC@Z was 3.75 × 10^-^3 cm/s (Supplementary Fig. **5a-f**), suggesting a slight decrease in the electron transfer rate of C in a bifunctionalised system.

**Figure 2.**
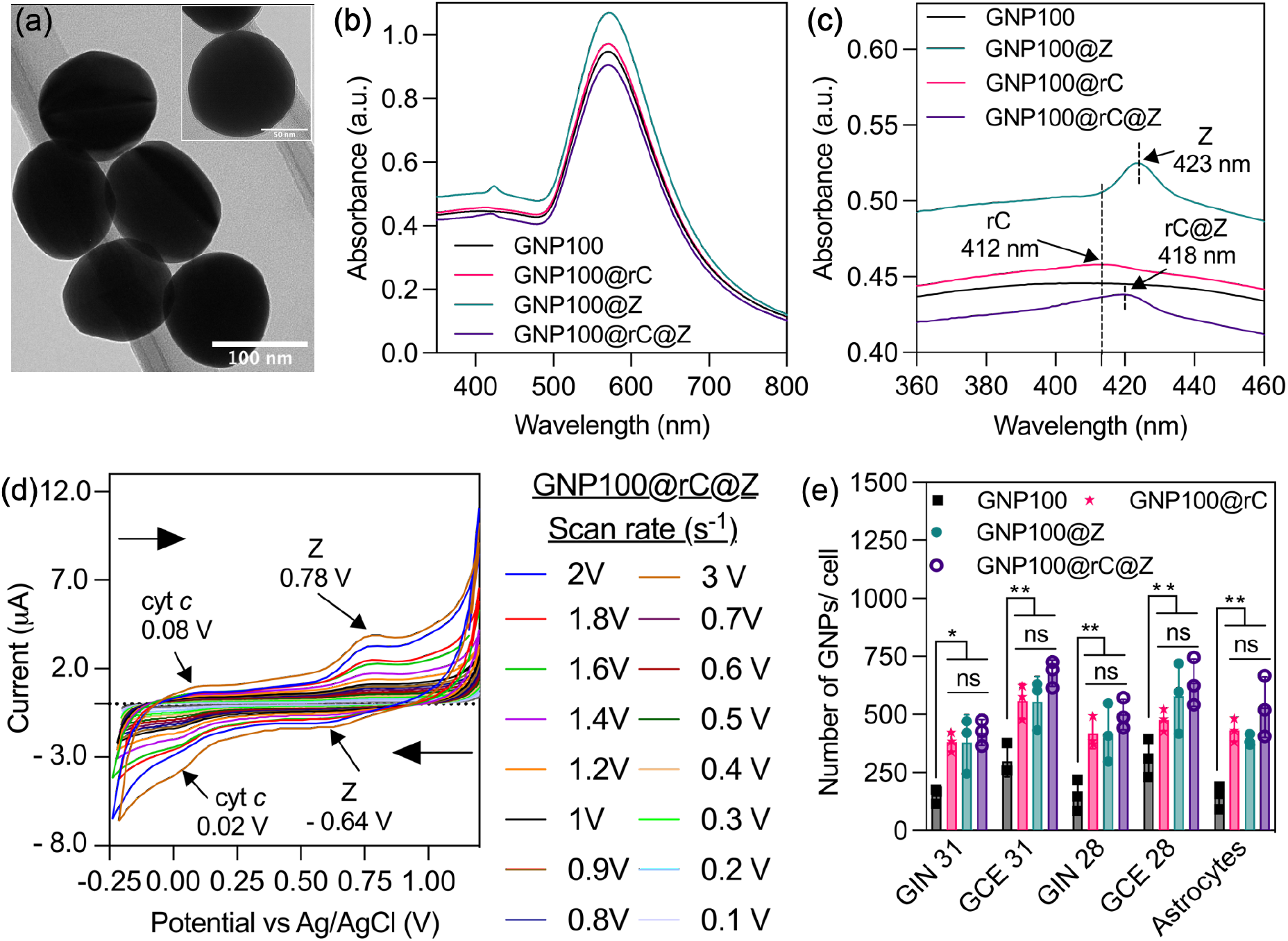
Physico-chemical & electro-analytical characterisation of bio-nanoantennae and their interaction with patient derived GBM cells. **(a)** Transmission electron microscopy (TEM) of bio-nanoantennae (GNP100@rC@Z) prepared by coupling reduced C (rC) and Zinc porphyrin (Z) to 100 nm PEGylated gold nanoparticles (GNP100). Inset represent HR-TEM image of 100 nm bio-nanoantennae. **(b)** UV-Vis absorption spectrum of bio-nanoantennae dispersed in PBS before ES. **(c)** Zoom-in UV-Vis absorption spectrum from 360-460 nm showing depict surface functionalisation with rC (GNP100@rC), Z (GNP100@Z), and both rC & Z (GNP100@rC@Z). **(d)** Cyclic voltammetry scan rate studies to analyse the redox properties of GNP100@rC@Z. Redox potentials were measured using an Indium tin oxide (ITO) working electrode, platinum wire counter and Ag/AgCl reference electrode with samples (25 µg/mL) dispersed in 10 mM PBS, scanned from +1.2 V to -0.25 V. **(e)** Inductively coupled plasma – mass spectrometry (ICP-MS) analysis to determine the association of bio-nanoantennae with different patient derived GBM cells and cortical astrocytes, the data is expressed as the number of GNPs per cell. Results are expressed as mean – S.D. \**p* < 0.05; \*\*\*\**p* < 0.0001 vs GNP100, obtained using 2-way ANOVA with a Tukey post-test.

To explore the potential of bio-nanoantennae on AC EF mediated redox switching of C, we used four types of preclinical GBM cells, isolated from two GBM patients: Glioma Invasive Margin (GIN 28 and GIN 31, isolated from an infiltrative tumor), Glioma Contrast-Enhanced core (GCE 28 and GCE 31, isolated from central tumor), which echo similar characteristics to different regions of GBM tumours^27^ and a commercial GBM cancer cell line U251. Human derived cortical astrocytes were also included as a control for non-tumorigenic cells. 3D analysis of z stacks confocal microscopy images (Supplementary Fig. 6) confirmed that the particles were internalised by all cell lines used after 8 hours of incubation, which were quantified using ICP-MS (Fig. **2e**). PrestoBlue assay data revealed that the particles are biocompatible up to a tested range of 100 µg/mL (Supplementary Fig. **7**).

## AC EFs mediated intracellular wireless electrochemistry on bio-nanoantennae surface induces apoptosis in pre-clinical GBM models

To examine the electrical input that can be sensed by intracellular bio-nanoantennae for inducing wireless electrochemistry, we optimised the voltage and frequency (Supplementary Fig. **8**) by assessing the change in metabolic activity of GIN 31 cells as the preliminary readout of C redox switching. The maximum effect on the metabolic activity was observed at 3 MHz, at an applied potential of 1V/cm and 0.65V/cm (no significant difference between 0.65V/cm *vs* 1V/cm, *p* value = 0.23) with significant differences between the tested controls (GNP100, GNP100@rC, and GNP100@Z) and the bifunctionalised bio-nanoantennae (GNP100@rC@Z). Importantly, the observed decrease of metabolic activity of GNP100@rC@Z treated cells was significantly higher than that following 24-hour application of FDA approved Tumour treating Fields (TTFs) *in vitro*.^28,29^ We ascribe this significant decrease in metabolic activity to the electrical-molecular communication via redox switching of rC to oxidised form of C (oC), thus inducing cell stress. To eliminate any potential effect of applied 1 V on water electrolysis (1.23 V *vs* normal hydrogen electrode), we chose 0.65 V/cm for further studies. The response of the cells to the treatment with GNP100@rC@Z bio-nanoantennae was comparable in different patient derived GIN and GCE cells, as well as in U251 cell lines (Fig. **3a-b** and Supplementary **Fig 9**), with a significant ∼ 50% decrease in metabolic activity achieved after 12 h *vs* a ∼ 20% decrease after 2 h AC EF stimulation (Supplementary Fig. **10**). This decrease in metabolic activity was significantly higher compared to the control (*p* values obtained from the statistical analysis are listed in supplementary table **6**). On the other hand, a significantly weaker effect (∼ 20% decrease in metabolic activity) was observed in cortical astrocytes, which was found to be not significantly different from the control (Fig. **3c**). Thus, based on the obtained data, we draw a conclusion that the change in metabolic activity depends on the duration of treatment and cell type.

**Figure 3.**
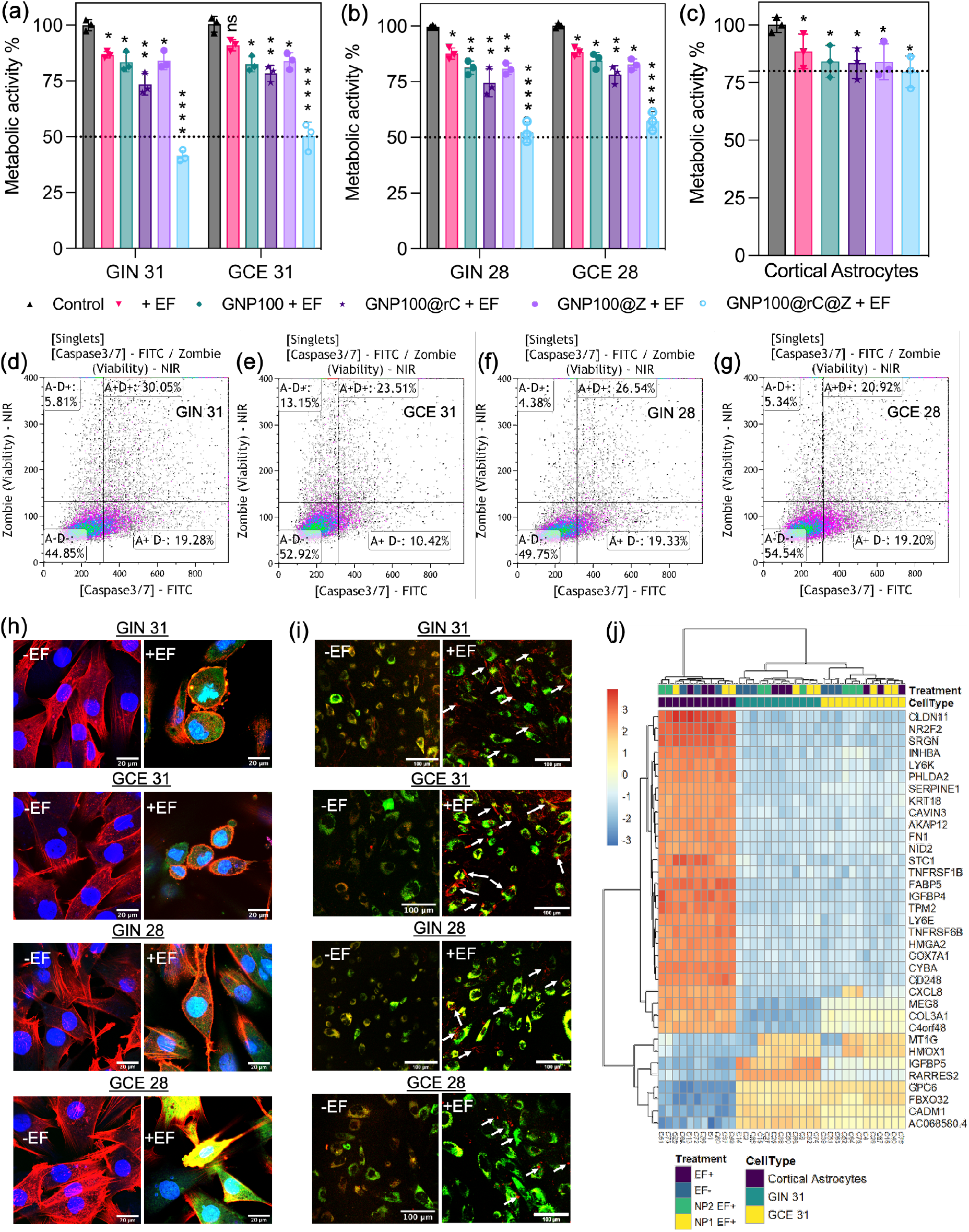
AC-EF responsive bio-nanoantennae mediated wireless electrical molecular communication induces caspase 3/7 mediated apoptosis in preclinical glioblastoma cells. **(a-c)** Metabolic activity of GIN/GCE 31, GIN/GCE 28, and cortical astrocytes was analysed using PrestoBlue HS assay. GIN, GCE, and human cortical astrocytes were treated with GNP100@rC@Z for 8 h followed by AC-EFs stimulation (3MHz, 0.65V/cm) for 12 h. Error bars represent mean ± standard error of mean (S.E.M.) obtained from triplicate experiments repeated thrice. Statistical analysis was performed by applying 2-way ANOVA with a Tukey’s post-test **(d-g)** Representative samples of flow cytometric analysis of cells stained with CellEvent Caspase-3/7 Green to detect caspase 3/7 apoptotic activity and Zombie NIR fixable dye to detect dead cell population. The quadrants in the figure represent the following: A+ = caspase 3/7 positive apoptotic cells and D+ = non-viable/dead cells. **(h)** High-magnification confocal microscopy images to demonstrate caspase 3/7 activation immediately after the treatment with AC-EFs (3MHz, 0.65V/cm) for 12 h in presence of bio-nanoantennae. Cells were fixed with paraformaldehyde followed by counterstaining with caspase 3/7 green detection kit, Cytopainter actin phalloidin (Texas Red 591 red), and Hoechst nuclear stain (blue). Scale bar = 20 mm. **(i)** Confocal microscopy image to demonstrate cytoplasmic localisation of GNP100@rC@Z immediately after the treatment with AC-EFs (3MHz, 0.65V/cm) for 12 h in presence of bio-nanoantennae. Cells were stained with late endosome dye (green) and imaged using a Leica confocal microscope with GFP (late endosomes) and Alexa 633 (GNP100@rC@Z) filter settings. Scale bar = 100 mm. **(j)** Heat map demonstrating hierarchical clustering of top 35 genes that were regulated after the treatment with GNP100@rC@Z for 8 h followed by AC-EFs stimulation (3MHz, 0.65V/cm) for 2-hour. A variance stabilised transformation was performed on the raw count matrix and 35 genes with the highest variance across samples were selected for hierarchical clustering. Each row represents one gene, and each column represents one sample. The colour represents the difference between the count value to the row mean. GIN/GCE 31 which showed maximum response to the treatment with GNP100@rC@Z and AC EFs were chosen. As a control for cancer cells, healthy cortical astrocytes were used. Immediately after the treatment, cells were washed and centrifuged to obtain a pellet, which was snap frozen in liquid nitrogen and shipped (in dry ice) to Qiagen, Ltd, Germany for RNA sequencing. The treatment codes in Fig. 1j are as follows: - EF = Control (no treatment with either bio-nanoantennae or AC EFs); + EF = cells treated with AC EFs; NP1 EF+ = cell treated with GNP100@rC, and NP2 EF+ = cells treated with GNP100@rC@Z, for 8 h followed by 2-hour AC EFs.

The alteration in metabolic activity in cells treated with GNP100@rC@Z + AC EFs (3 MHz at 0.65 V/cm for 12 hours) is correlated with increased cell death, as clearly evidenced by the results of live-dead assay (Supplementary **Figs. 11-12**). The mechanism of cell death was probed by flow cytometry (Fig. **3d-g** and Supplementary Figs. **13-14**) and confocal microscopy (Fig. **3h** and Supplementary Fig. **15**), where the induced caspase 3/7 activity was observed in GIN/GCE cells treated with GNP@rC@Z followed by ES, indicative of apoptosis.^30^ confocal microscopy revealed that bio-nanoantennae are localised within the cytosol (Fig. **3i**). We note that no reactive oxygen species and temperature changes were observed (Supplementary Figs. **16-17**). Therefore, we successfully conducted ES of GBM cells and demonstrated wireless electrochemistry mediated redox switching and activation of C on the surface of GNP100@rC@Z leading to apoptosis of GBM cells.

To further understand this electrical-molecular signalling. We performed transcriptomic analysis on a sample of glioblastoma cells and astrocytes (Fig **3j** and Supplementary Figs. **S18-19)** to interrogate the effect of the bioelectronic communication tool on gene expression and regulation. Hierarchical clustering analysis showed differential expression of genes related to apoptosis, cancer proliferation and angiogenesis, and tumour suppression. This change in differentially expressed genes was highest for GIN/GCE 31 cells treated with GNP100@rC@Z followed by ES compared to untreated cells and other experimental controls. Gene ontology analysis revealed that most of the differentially upregulated genes such as CXCL8 & INHBA in GCE 31, and MT1G & HMOX1 in GIN/GCE 31 are related to apoptosis, implying that apoptosis was upregulated after the treatment.^30–33^ Furthermore, the upregulation of HMOX1 (a haem oxygenase) could also be a result of the presence of haem containing cyt *c* and Z on bio-nanoantennae. On the other hand, most downregulated genes such as STC1 (GIN 31), IGFBP5 (GIN/GCE 31), and FBXO32 (GIN/GCE 31) are characteristic of angiogenesis in cancer proliferation, and tumour growth and metastasis, respectively.^34–37^ Interestingly, we did not observe any changes in the gene regulation for astrocytes, which suggests that the treatment modulates signalling pathways that are specific only to GBM. Further studies are underway to ascertain the cell signalling pathways that are implicated during this treatment. Altogether the obtained data implies that GNP100@rC@Z treatment followed by ES leads to an increase in apoptosis, and reduced proliferation and invasiveness of patient derived GBM cells. Overall, the *in vitro* results indicate successful communication with biology at a molecular scale using electricity. Importantly, this approach enables selective actuation of cancer cell behaviour (diseased *vs* healthy cells) compared to astrocytes.

## Nanoscale wireless electrochemistry and in vitro QBET

The electrical-molecular communication via wireless electrochemistry induced at GNP100@rC@Z, was probed by circular dichroism (CD) analysis (Fig. **4a** and Supplementary Fig. **20a-e**). The soret CD band of GNP100@rC@Z revealed two positive maxima (407 nm and 434 nm) suggesting the oxidation of the haem centre of C after ES with AC EFs of 3 MHz at 0.65 V/cm. This effect was corroborated by comparison with CD of native oC. Furthermore, one negative minimum (417 nm) was observed for native oC, which was red shifted to 419 nm indicating slight perturbation around the haem moiety. On the other hand, when a control experiment was carried out by performing ES of GNP100@rC@Z using insulated electrodes, we did not observe any signals from rC or oC. To further ascertain nanoscale BPE, the UV-Vis spectrum of GNP100@rC@Z was monitored after the ES (Fig. **4b-c** and Supplementary Fig. **21 a-c**), and revealed a significant blue shift in the absorbance peak of rC from 412 nm to 408 nm and red shift in the absorbance peak of Z from 423 nm to 432 nm, which can be attributed to the oxidation of the haem moieties in C and a change in the environment of Z leading to the formation of J aggregates,^25^ respectively. We note that no shift of absorption peaks was observed without EF stimulation (Supplementary Fig. **22).** The obtained spectroscopic data confirms that we were successfully able to induce nanoscale BPE on the surface of GNP100@rC@Z to modulate the redox state of C (Fe^2+^ to Fe^3+^) using remotely controlled AC EFs. In addition, there was no significant change in *h*_d_ and ζ after electrical stimulation (ES) with AC electric fields (AC EF) of 3 MHz at 0.65V/cm suggesting the surface chemistry of the synthesised nanoantennae is stable (Supplementary Figs. **23-24**). Thus, we demonstrate that the bio-nanoantennae act as bipolar electrodes and can modulate the redox state of the C under application of the EF. We envisage that these processes can be used to provide molecular communication with the cells to modulate their function.

**Figure 4.**
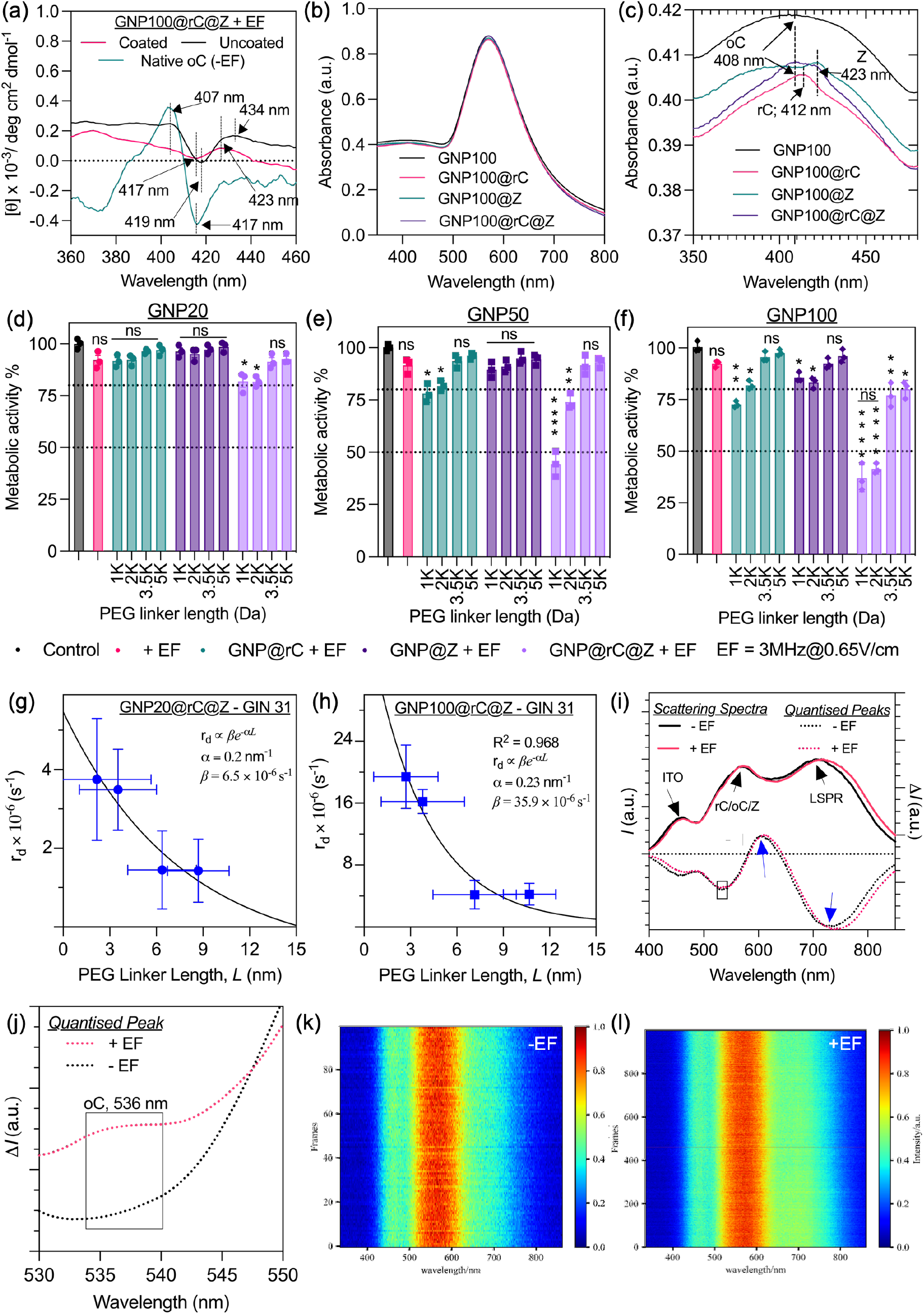
Nanoscale wireless electrochemistry and *in vitro* electron tunnelling via bio-nanoantennae for inducing cell death in GBM cells. **(a)** The effect of AC-EFs on the conformation of redox moieties: soret CD spectrum to emphasise redox mediated change in haem moieties of free native oC, rC, and bifunctionalized GNP nanoactuators in 10 mM PBS (pH = 7.4). All samples with identical concentrations (25 µg/mL) were used for spectrum acquisition. Three spectra of each sample were collected and averaged. Insulated coated steel electrodes were used as a control to demonstrate the occurrence of wireless electrochemistry at nanoscale. **(b & c)** UV-Vis spectrum of bio-nanoantennae after stimulation with AC-EFs of 3MHz, 0.65V/cm indicating blue shift in absorption maxima of rC to oC (oxidised C) in GNP100@rC@Z indicating redox switching of C. **(d-f)** Metabolic activity of GIN 31 cells as function of different size (20 nm, 50 nm, and 100 nm) bio-nanoantennae (GNP20@rC@Z, GNP50@rC@Z, and GNP100@rC@Z) synthesised using various linker lengths (1k, 2k, 3.5k, and 5k Da). GIN 31 cells were treated with bifunctionalised bio-nanoantennae for 8 h followed by AC-EFs stimulation (3 MHz, 0.65V/cm) for 12 h. **(g-h)** Mathematic modelling to determine the rate of donor charging, *r_d_*, calculated using the metabolic activity, using equation no. 1, compared to the PEG linker length, *L*, for (d) GNP100@rC@Z and (e) GNP20@rC@Z in GIN31 cell samples. The black lines show the exponential behaviour expected for quantum tunnelling with an inverse localisation radius *α* and constant of proportionality *β* shown in the figure legends. **(i)** Scattering spectra and spectra difference for QBET obtained for GNP100@rC@Z bio-nanoantennae. The quantised peaks were obtained from the difference of scattering spectra between the samples functionalised with rC and Z using 2000 Da linker and GNP100. Solid curves are captured scattering spectra (linked to left axis) of GNP100@rC@Z, and dashed curves are quantised peaks i.e., the corresponding spectra difference (linked to right axis). Blue boxes indicate peak shift. **(j)** Quantised peaks GNP100@rC@Z within in 530-550 nm region confirming the presence of oC in samples exposed to AC EFs. **(k-l)** Two-dimensional heat map of scattering spectral changes obtained from dark field microscopy, in the presence and absence of EF.

To facilitate the translation of the technology more broadly, understanding of the electrically induced mode of electron transfer is needed. The frequency we used for electrical communication was 3 MHz Considering the electron transfer rate constant of 3.75 × 10^-^3 cm/s calculated from Fig **2d**, the maximum distance the electron could travel at this frequency is 0.01 nm. Thus, theoretically at this given frequency, the redox event should not occur. We propose that the observed activity is enabled by QBET induced from the *C* as was indicated previously.^10^ However this process has never been controlled through electrical stimulation.

To gather experimental evidence for QBET in our system, we consider molecular tunnel junctions (MTJs), where quantum electron tunnelling can occur.^39–42^ Therefore, we altered the tunnel junction energy by using GNP with different sizes (20, 50, and 100 nm), and PEG linker with different lengths (1, 2, 3.5 and 5k Da) (Supplementary Figs. **S25**-**26**). The metabolic assays (Fig **4d-f** & Supplementary Figs. **27-28**) and viability studies (Supplementary Figs. **29-30**) on GIN 31/GCE 31 cells with ES (3 MHz and 0.65V/cm) indicated a resonant biological effect with bio-nanoantennae using the 1 kDa PEG linker with 50 nm and 100 nm GNPs. Importantly, this effect is only seen for cells treated with bio-nanoantennae (GNP@rC@Z) following 12 h ES. ICP-MS analysis revealed no significant difference in the number of Z/rC per cell when treated with GNP20@rC@Z/ GNP50@rC@Z/ GNP100@rC@z (Supplementary Figs. **31-32**). The explanation for this result is garnered from the work of Hongbao Xin et al. who showed that the QBET could be captured from *C* by tuning the tunnel junction via the size of SAM linker.^10^ We envisage that a similar phenomenon is occurring in our system, as evidenced by the resonance at specific conditions and is indicative of wave-like behaviour.^43^ Thus, we suggest that the electron transfer is a QBET, and this is the first example of an electrically stimulated QBET in biology to date and consequently facilitating electrical-molecular communication.

To provide further evidence of the mechanism of electron transfer we established a mathematical model. The cell metabolic rate (Fig. **4d and 4f**) was correlated to the rate of charge transfer from the C derived equation no. 1 (full details of the derivation are presented in Supplementary information section 1.6. of materials and methods). These values were then plotted as a function of the barrier junction altered using the ligand for different sized nanoparticles (Fig. **4 g-h**).

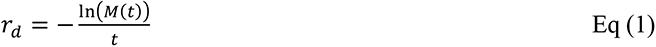

where *r*_d_ is rate of donor charging and *M*(*t*) is metabolic activity at time *t*.

The results are well fitted with exponential dependences expected for the tunnelling of an electron through a barrier,^44^ supporting the hypothesis that the resonance biological effects are observed with the electrical input results in a QBET. Further primary experimental evidence for the quantum tunnelling is provided by the PRS probing plasmon resonance energy transfer (PRET).^45^ Scattering spectra were obtained for GNP100@rC@Z with and without ES. Without ES, quantised dips were obtained by subtracting the spectra for the unmodified particle and vice-versa. For GNP100@rC@Z samples (no ES) the scattering peaks were observed at ∼ 465 nm corresponding to ITO, a broad peak due to rC and Z at around 568 nm and localised surface plasmon resonance (LSPR) peak shifted to 711 nm (GNP100 sample LSPR = 612 nm) (Fig. **4i** and Supplementary Fig **33a**). This equates to the frequencies with the absorption peak of rC/Z and subsequently induces the electronic excitation of the rC at the specific wavelength.^10^ Importantly, the QBETs captured are cumulative. Under ES, we see a quantised dip at 536 nm due to the switching of rC to oC and thus representing QBET (Fig. **4j**). We also see a 20 mV shift in the SPR peak (heat maps Fig. **4k**: no EF, and Fig. **4l**: EF applied) indicative of charge transfer, which we associate with the electron relay donor complex redox behaviour.^46,47^ It is important to note that the appearance of oC was not observed in all other control samples with ES (3MHz, 0.65V) (Supplementary Fig **33 b-i**). The data combined indicated resonant frequency resulting in cell death, model exponential decay based on barrier junction energetics and PRET data resulting in quantised dips that represent electron tunnelling. Therefore, to the best of our knowledge, this is the first successful demonstration of electrical-molecular quantum signalling technology in biology, which we demonstrated could be used for cancer cell induced apoptosis.

## Conclusions

The bio-nanoantennae system described here, based on gold nanoparticles functionalised with electron relay species and inspired by wireless electrochemistry, is sensitive to electric fields: The redox state of molecules functionalised on the surface of nanoantennae can be tuned. Electrical-molecular communication within glioblastoma cells can be performed using these antennae with an electric field. This effect was demonstrated through the stimulation of specific cell response in the form of apoptosis, achieved by modulating the redox state of the *cyt c*. We believe that the electron transfer in the bio-nanoantennae occurs from electric field-induced quantum tunnelling. Moreover, transcriptomics shows that electrical-molecular communication is targeted in cancer cells. This represents a wireless electrical-molecular communication tool that facilitates cancer cell killing.

## Acknowledgements

This work was supported by the Engineering and Physical Sciences Research Council Grant number [EP/R004072/1], Royal Society of Chemistry Enablement Grant E21-1135058786, University of Nottingham internal funding schemes - NanoPrime and UNICAS. The authors would like to thank Mr Baokang Xie at Nanjing University, China for PRS and DFM analysis, David Onion at University of Nottingham, Flow cytometry Unit for assistance with flow cytometry analysis, Dr Mike Fay for help with TEM, and Jordan Potts & Ruby Brown for assistance with particle analysis.

## 1. Materials and Methods

### 1.1. Synthesis of bifunctionalised gold bipolar bio-nanoantennae

Spherical gold nanoparticles (GNPs) with diameter (d) = 100 nm and capped with thiol-carboxylic-polyethylene glycol (SH-PEG-COOH; molecular weight *M* = 1KDa, 2KDa, 3KDa, and 5KDa) with PEG density of 1.5 nm^2^ were purchased from Nanopartz Inc. USA. The reduced cyt *c (rC) was obtained by adding 10 mg of* oxidised form of horse heart cyt *c* (oC) into 5 mL L-ascorbic acid solution (1 mg/mL in PBS) and purified by dialysis at temperature *T* = 4°C under dark condition for 36 h to remove excess ascorbic acid. The rC and Zinc 5-(4-aminophenyl)-10,15,20-(tri-4-sulfonatophenyl)porphyrin triammonium (Z) were covalently conjugated to carboxylic group on capping ligands of GNPs using EDC/NHS carbodiimide coupling chemistry. Briefly, 20 μL of GNP solution (3.6 mg/mL in ultrapure water) were mixed with 20 μL of EDC/NHS solution in 2-(N-morpholino)ethane sulfonic acid (MES) buffer (10 mM, pH = 5.5) at a concentration of 30 and 36 mg/mL. The solution was mixed for 1h at room temperature, then 1 mL of washing buffer (1X phosphate buffer saline (PBS) with 0.01 % w/v Tween 20) was added, and the solution was centrifuged at 2500 rpm for 20 min. The supernatant was discarded and 20 μL of rC (1 mg/mL) and Z (0.5 mg/mL) was added to the pellet and sonicated using a Fisherbrand ultrasonic bath (FB11201, 80 KHz, 50% power, 1 min). Next, the solution was incubated for 4 h at room temperature under mixing, then 1 mL of washing buffer was added, and the solution was centrifuged at 2500 rpm for 20 min. To ensure, complete removal of unbound rC and Z the washing step was repeated twice. The obtained pellet rC and Z conjugated GNPs (GNP100@rC@Z) was dispersed in PBS and stored at 4 °C until further use.

The control samples of gold nanoparticles covalently conjugated with only one molecule, either rC (GNP100@rC) or Z (GNP100@Z), only one of these compounds was introduced during EDC/NHS step with concentrations of 0.25 mg/mL and 0.1 mg/mL, respectively. The same protocol was used to synthesise bio-nanoantennae with different GNP diameters (20 nm, 50 nm, and 100 nm) and different PEG length (1 kDa, 2 kDa, 5 kDa). The concentration of cyt *c* and Z during EDC/NHS step was optimised (0.25, 0.5, and 1 mg/mL) to obtain similar binding ratio of cyt *c*/Z per GNP.

### 1.2. Characterization Techniques

#### 1.2.1. Dynamic light scattering (DLS) and Zeta potential measurements

The hydrodynamic diameter (h_d_) and Zeta potential (ζ) of bio-nanoantennae (in ultrapure water) was measured using a Malvern Zetasizer Nano-ZS ((Malvern Instruments, UK).

#### 1.2.2. Transmission electron microscopy (TEM)

Transmission electron microscope (JEOL 2000 FX TEM) operating at 200 kV accelerating voltage was used to record TEM images. The samples were prepared by drop-casting 10 μL of sample onto a carbon-coated copper grid (400 Mesh, Agar Scientific) twice in an interval of 1 h. The sample was dried for at least 30 min before TEM imaging.

#### 1.2.3. UV-Vis absorption spectroscopy

UV–Vis absorption spectra of bio-nanoantennae dispersed in PBS were recorded on a Cary 3500 UV-Vis (Agilent Technologies Ltd).

#### 1.2.4. Circular dichroism (CD)

Far and near UV CD spectra were recorded at *T* = 20°C on a Chirascan CD spectrophotometer (Applied Photophysics) equipped with a temperature control unit TC125 (Quantum Northwest). Samples were dispersed in 10 mM PBS at pH 7.4. At least 3 spectra were recorded for each sample and averaged. A quartz cuvette with an optical pathlength of 1 cm was used for the CD measurements.

#### 1.2.5. Cyclic voltammetry (CV)

The electrochemical analyses were conducted using a Metrohm Autolab M204 potentiostat equipped with the three-electrode system consisting of a platinum wire counter electrode, an Ag/AgCl reference electrode (ALS Co. Ltd.) and an indium tin oxide (ITO, Delta Technologies Ltd.) as working electrode. ITO coated glass (10 mm × 20 mm) was washed with acetone and water, dried with argon, and assembled into an electrochemical cell with an exposed working area of 38.5 mm^2^. Bifunctionalized gold bipolar nanoelectrodes were dispersed in PBS to a final concentration of 25 μg/mL (determined using UV–Vis). CV was conducted between 1.2 V and - 0.2 V with varying scan rates between 50 mV·s^-1^ and 2 V·s^-1^. Repetitive consecutive CVs were conducted at a fixed scan rate of 100 mV·s^-1^. Control CVs were conducted with carboxylic PEG modified gold nanoparticles using PBS as the supporting electrolyte.

Heterogenous rate constant (*k°*), was calculated using Nicholson and Shain method.^1^ The rate transfer coefficient, α, was calculated from the scan rate study (Supplementary Fig **5**). The slope of the logarithm of the scan rate versus the difference between the peak potential and formal potential of the cell is given by the Equation (1):

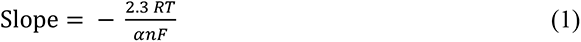

Where *R* is gas constant; and *T* is temperature, *n* is number of electrons transferred in the redox reaction *F* is Faraday constant.

However, Nicholson and Shain method for determining *ψ* assumes that *α* = 0.5. However, in this work the value *α* is different (Supplementary, Table **5**), therefore Lavagnini method^2^ is used to calculate *ψ* as the function of the peak separation Δ*Ep* using Equation (2):

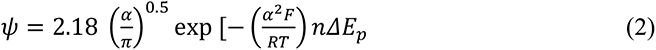

Then *k°* for the nanoantennae is calculated using the Equation (3):

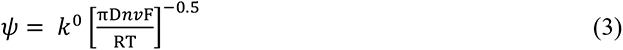

where *D* is diffusion coefficient.

### 1.3. Cell culture studies

#### 1.3.1. Cell lines

Glioma INvasive Marginal (GIN) cells were isolated from the 5-aminolevulinic acid (5-ALA) fluorescing infiltrative tumour margin and Glioma Core Enhanced (GCE) cells were isolated from the core central region of the tumor from the glioblastoma (GBM) patients, who underwent surgery at the Queen’s Medical Centre, University of Nottingham (Nottingham, UK) using previously described method.^3^ Low-passage U251 cell lines (purchased from ATCC, USA) and patient-derived (GIN 28, GIN 31, GCE 28, and GCE 31 cells were cultured in DMEM (Gibco) supplemented with 10% FBS, 1% Penicillin/Streptomycin and 1% L-Glutamine. Human derived cortical astrocytes (HCA) (Cat. No. 1800, Batch No. 24490, ScienCell) were cultured in astrocyte medium (AM) containing 2% FBS, 1% astrocyte growth supplement, 1% Penicillin/Streptomycin from ScienCell. All cells were maintained at *T* = 37°C in an incubator with humidified atmosphere, containing 5% CO_2_. Cells were routinely tested for mycoplasma (once a month) where they were grown in an antibiotic-free medium for one week before mycoplasma testing. All cells used were mycoplasma-free.

#### 1.3.2. PrestoBlue HS assay for biocompatibility and metabolic activity studies

The cells (U251, HCA, GIN and GCE) were seeded in a 96-well at a density of 5 × 10^3^ cells/well and allowed to adhere for 24 h. The media was replaced with fresh media containing GNP conjugates at different concentrations (25, 50, and 100 μg/mL) and the cells were incubated for 8 h. The media was removed, and the cells were washed with PBS and incubated for another 48 h in fresh media. The media was replaced with complete media containing 10% PrestoBlue^TM^ HS cell viability reagent (ThermoFisher Scientific, UK) and incubated for an hour before reading the fluorescence at 590 nm / 610 nm (excitation/ emission) in a Tecan microplate reader (Infinite M Plex and Spark 10M). Cells grown in culture media only provided the negative control. Values are presented relative to negative control. The data is represented as an average of triplicate experiment with 3 independent repeats.

#### 1.3.3. Cellular association and uptake

GIN and GCE cells were seeded into a 24-well plate at density of 1 × 10^5^ cells per well and incubated at 37 °C for 24 h. After 24 h, the culture medium was replaced with fresh medium containing 25 μg/mL of GNP100@rC@Z and incubated for 8 h. Then the media was removed, and the cells were washed with PBS (300 μL, repeated 2-times). The cells were trypsinised and 50 μL of cell suspension was used for trypan blue cell viability and counting. The remaining cell suspension was centrifuged at 300 g for 5 min. The obtain cell pellet was digested overnight with 70% nitric acid, diluted with milliQ water to reach the acid concentration to 2% and taken for ICP-MS analysis (iCAPQ Thermo Fischer). To confirm the cellular uptake of bipolar nanoelectrode, after 8 h exposure, the cells were washed with PBS (300 μL, two times), fixed with 4% paraformaldehyde for 15 min and washed twice with PBS. The cell nuclei were stained with Hoechst 33342 (NucBlue™ Live Ready Probes™ Reagent, ThermoFisher Scientific, UK) and actin using Phalloidin-iFluor™ 488 conjugate (AAT Bioquest, Stratech, UK) and incubated for 1 h at 37°C in the dark. After washing the cells twice and immersing them in PBS, the fluorescence imaging was performed using Leica TCS SPE Confocal Microscope. The orthogonal sections of z-stacks were obtained, and the images were analysed using ImageJ.

### 1.4. Electrical stimulation (ES) studies

#### 1.4.1. Electrical stimulation (ES) of cells

U251, GIN 28, GIN 31, GCE 28, and GCE 31 cells were seeded in a 24-well plate (μ-plate 24 well black, ibiTreat, Thistle Scientific, UK) at a density of 7.5 × 10^4^ cells/well, while HCA were seeded at a density of 5 × 10^4^ cells/well in a Poly-L-Lysine coated 24-well plate. The cells were incubated for 24 h at 37°C and 5% CO_2_, then the cell culture medium was replaced with fresh medium containing bio-nanoantennae (25 μg/mL) and incubated for 8 h. The cells were washed twice with PBS. Two steel electrodes (0.5 mm × 25 mm) were placed at a fixed distance (at opposite sides of the well and 10 mm from each other) into each well of a 24-well plate and dipped in cell culture media. The electrodes were connected to an Arbitrary Function Generator (AFG-21225, RS PRO, UK) to deliver the required AC sine-wave signals, frequency, and amplitude. The cells were stimulated with alternating current electric fields (AC-EFs) with frequency of 3 MHz and a peak voltage amplitude of 0.65 V/cm for a period of 2 h or 12 h. The EF between the electrodes was measured using a digital oscilloscope (TDS 210, Tektronix) and the temperature was monitored every 2 h using an InfraRed laser gun (IR-801, ATP). The intensity of EF is expressed in peak voltage amplitude per centimetre (V/cm). The metabolic activity of cells after electrical stimulation was analysed using PrestoBlue HS assay (Section 1.3.2).

#### 1.4.2. Calcein AM, and Propidium Iodide (PI) live/dead assay

Immediately after the ES, the media was removed and replaced with fresh media containing mixed dyes 1μM Calcein AM and 1μg/mL PI (ThermoFisher, UK), and incubated for 30 min at 37°C and 5% CO_2_. The cells were washed twice with PBS and fresh phenol red free medium was added. The cells were imaged using a Nikon Eclipse Ti fluorescent microscope with GFP and mCherry filter settings. The population of live and dead cells were quantified using ImageJ software.

#### 1.4.3. H_2_DCFDA/DCF reactive oxygen species (ROS) generation assay

The cells were incubated with nonfluorescent cell-permeant 2’,7’-dichlorodihydrofluorescein diacetate (H_2_DCFDA, 5μM, ThermoFisher Scientific, UK) probe for 30 min prior to ES. Immediately after the ES, the cells were washed with PBS. The generated ROS converted H_2_DCFDA into 2’,7’-dichlorofluorescein (DCF). Green fluorescence of DCF was detected using Nikon Eclipse Ti with FITC filter settings.

#### 1.4.4. Caspase 3/7 flow cytometry analysis of cell death

Immediately after the ES, cells were trypsinised and centrifuged (300g for 5 min) to obtain cell pellet. After washing with PBS, cells were incubated with a dye master mix containing CellEvent^TM^ Caspase 3/7 Green Detection Reagent (ThermoFisher Scientific, 1:1000) and Zombie NIR fixable viability stain (BioLegend, 1: 2500) for 30 min. Then the cells were centrifuged at 300 g for 5 min, washed with PBS and fixed with 4% paraformaldehyde. The fluorescence signal of Caspase 3/7 (Excitation / Emission = 511 nm / 523 nm) and Zombie NIR dye (Excitation / Emission = 719 nm / 746 nm), characteristic for apoptotic and necrotic cell population, respectively, was detected using ID7000 Spectral Flow Cytometer. Kaluza software (v2.1) was used to analyse the data.

#### 1.4.5. Caspase 3/7 Detection using confocal microscope

Immediately after the ES, the media was removed and the cells were incubated with 8 µM CellEventTM Caspase 3-7 green detection reagent (Thermo Fisher) in PBS containing 5% FBS for 30 min at 37 °C. Afterwards, the cells were fixed with 4% paraformaldehyde for 20 min and subsequently washed twice with PBS. Later the cells were treated actin stain Phalloidin-iFluor™ 594 conjugate (AAT Bioquest, Stratech, UK) for 90 min at 37°C in the dark and washed again with PBS followed by staining with Hoechst 33342 for 10 min. Finally, the cells were washed with PBS (twice) and imaged using a Leica TCS SPE Confocal Microscope at 63x objective using the filter settings of Alexa Fluor 488 and Texas Red 594 dye.

#### 1.4.6. Co-localisation studies

GIN and GCE cells were seeded at a density of 4 × 10^4^ cells/ well in a 24-well plate and incubated at 37 °C for 24 h. After 24 h, the culture medium was replaced with fresh medium containing CellLight™ Late Endosomes-GFP, BacMam 2.0 (ThermoFisher Scientific, UK) and incubated overnight at 37 °C and 5% CO_2_. Later, the media was replaced with fresh media containing 25 μg/mL of GNP100@rC@Z and incubated for 8 h. Immediately after the ES, the cells were washed with PBS and imaged using Leica confocal microscope.

### 1.5. Gene regulation analysis

#### 1.5.1. Differential gene regulation analysis

Immediately after the ES (3 MHz, 0.65V/cm, 2 hours) cells were washed with PBS, trypsinised, and centrifuged to obtain a pellet. The cell pellets were snap frozen in liquid nitrogen for 5 min and stored at *T* = - 80°C until shipment (in dry ice) to Qiagen genomics facility at Hilden, Germany.

#### 1.5.2. Sample preparation

RNA was isolated from 200000 cells using the RNeasy Micro (QIAGEN) according to manufacturer’s instructions with an elution volume of 14µL.

#### 1.5.3. Library preparation and sequencing

The library preparation was done using the QIAseq UPX 3’ Transcriptome Kit (QIAGEN). A total of 10ng purified RNA was converted into cDNA NGS libraries. During reverse transcription, each cell is tagged with a unique ID and each RNA molecule is tagged with a unique molecular index (UMI). Then RNA was converted to cDNA. The cDNA was amplified, during the PCR indices were added and the libraries were purified. Library preparation was quality controlled using capillary electrophoresis (Agilent DNA 7500 Chip). Based on quality of the inserts and the concentration measurements the libraries were pooled in equimolar ratios. The library pool(s) were quantified using qPCR. The library pool was then sequenced on a NextSeq (Illumina Inc.) sequencing instrument according to the manufacturer instructions with 100bp read length for read 1 and 27bp for read2. Raw data was de-multiplexed and FASTQ files for each sample were generated using the bcl2fastq2 software (Illumina inc.).

#### 1.5.4. Read demultiplexing, mapping, and quantification of gene expression

The “Demultiplex QIAseq UPX 3’ reads” tool of the CLC Genomics Workbench 20.0.4 was used to demultiplex the raw sequencing reads according to the sample indices. The “Quantify QIAseq UPX 3’ workflow” was used to process the demultiplexed sequencing reads with default settings. In short, the reads are annotated with their UMI and are then trimmed for poly(A) and adapter sequences, minimum reads length (15 nucleotides), read quality, and ambiguous nucleotides (maximum of 2). They are then deduplicated using their UMI. Reads are grouped into UMI groups when they (1) start at the same position based on the end of the read to which the UMI is ligated (i.e., Read2 for paired data), (2) are from the same strand, and (3) have identical UMIs. Groups that contain only one read (singletons) are merged into non-singleton groups if the singleton’s UMI can be converted to a UMI of a non-singleton group by introducing an SNP (the biggest group is chosen). The reads were then mapped to the Human genome hg38 and annotated using the refseq GRCh38.p13 mRNA annotation.

The ‘Empirical analysis of DGE’ algorithm of the CLC Genomics Workbench 21.0.4 was used for differential expression analysis with default settings. It is an implementation of the ‘Exact Test’ for two-group comparisons developed by Robinson and Smyth, 2008 and incorporated in the EdgeR Bioconductor package Robinson et al., 2010.

For all unsupervised analysis, only genes were considered with at least 10 counts summed over all samples. A variance stabilizing transformation was performed on the raw count matrix using the function vst of the R package DESeq2 version 1.28.1. 500 genes with the highest variance were used for the principal component analysis. The variance was calculated agnostically to the pre-defined groups (blind=TRUE). 35 genes with the highest variance across samples were selected for hierarchical clustering.

### 1.6. Model of Metabolic Activity, Charging Rate, and Quantum Tunnelling

We looked to develop a mathematical model to support the characteristic exponential decay connected with quantum mechanics,

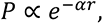

where *P* is the probability of electron tunnelling, *α* is the inverse localisation length, which scales with the energy barrier that an electron must tunnel through, and *r* is the length of the energy barrier. We define an intrinsic rate of cell death (directly proportional to probability of a given cell dying within a given time frame), *r_d_*, which we assume is proportional to the rate of cytochrome charging. Then, the number of dead cells (D) at time *t* is

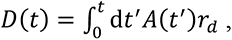

where *A* is the number of alive cells. The metabolic activity is given by

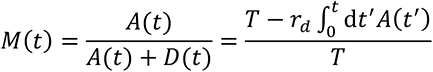

Where, *T*=*D*+*A* is the total number of cells. Solving the above equation, we find that *M*(*t*) = *e^−r_d_t^*. Therefore, the rate of charging at any given time is given by r_d_ α – ln*M*(*t*).

### 1.7. Dark-Field microscopy and plasmon resonance scattering (PRS) spectroscopy

The PRS measurements were carried out on an inverted dark-field microscope (eclipse Ti-U, Nikon, Japan) using a 40× objective lens (NA = 0.6) and a dark-field condenser (0.8 < NA < 0.95). A halogen lamp (100 W) was used as a source of white light to generate plasmon resonance scattering light. The dark-field images were captured by a true-color digital camera (Nikon DS-fi, Japan). The light scattered from the bifunctionalised nanoantennae was split by a monochromator (grating density: 300 lines/mm; blazed wavelength: 500 nm, Acton SP2300i, Princeton Instruments, USA). IsoPlane-320 spectrometer was used, and the split light was collected by a charge-coupled device (Pixis 100BX, Princeton Instruments, USA). Alternating current electric field of 3 MHz at 0.65 V were applied for 10 min and scattering spectra were monitored (1000 frames recorded). The exposure time was 500 ms. The samples for PRS were prepared by immobilising nanoantennae on ITO. Firstly, ITO slides were treatment with ethanol, acetone, and water under sonication. Next, 50 µL nanoantennae solution was drop casted on ITO slides for 10 min followed by a single step washing and rinsing with water. Finally, the slides were dried with N_2_ gas.

### 1.8. Statistics and reproducibility

All the statistical analyses were performed using GraphPad Prism v9.4.1 software (GraphPad Software, Inc). All the data is expressed as mean ± S.EM., unless specified. For responses that were affected by two variables, a 2-way ANOVA with a Tukey post-test was used. Results are expressed as mean ± standard deviation. The value P ≤ 0.05 was considered significant. The number of technical replicates and independent repeats are included in the figure legends.

## 2. Results and Discussions

**Figure 1.**
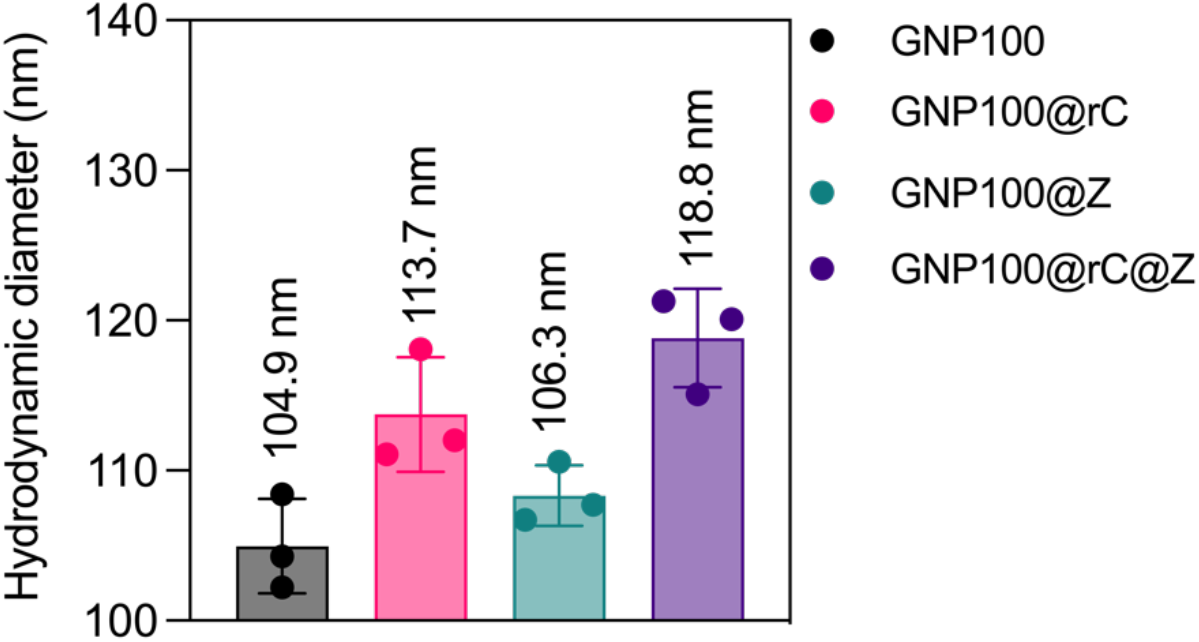
Hydrodynamic diameter of bio-nanoantennae (dispersed in ultra-pure water) before ES. Error bars represent mean ± s.d. obtained from 3 individual experiments.

**Figure 2.**
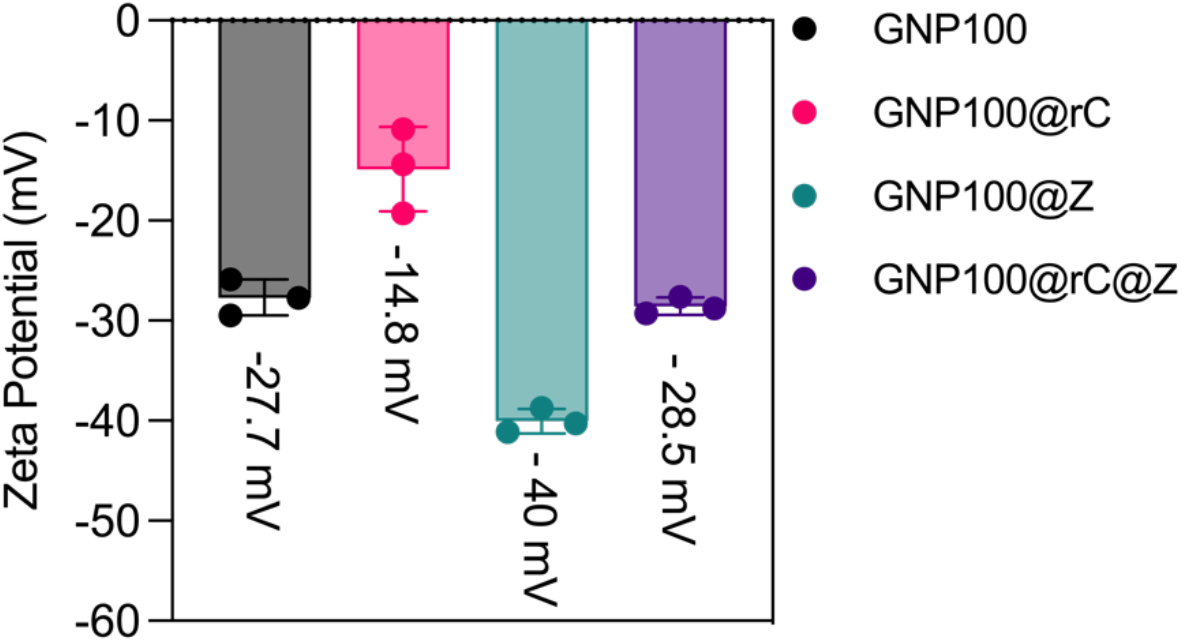
Zeta potential of bio-nanoantennae (dispersed in ultra-pure water) before ES. Error bars represent mean ± s.d. obtained from 3 individual experiments.

**Figure 3.**
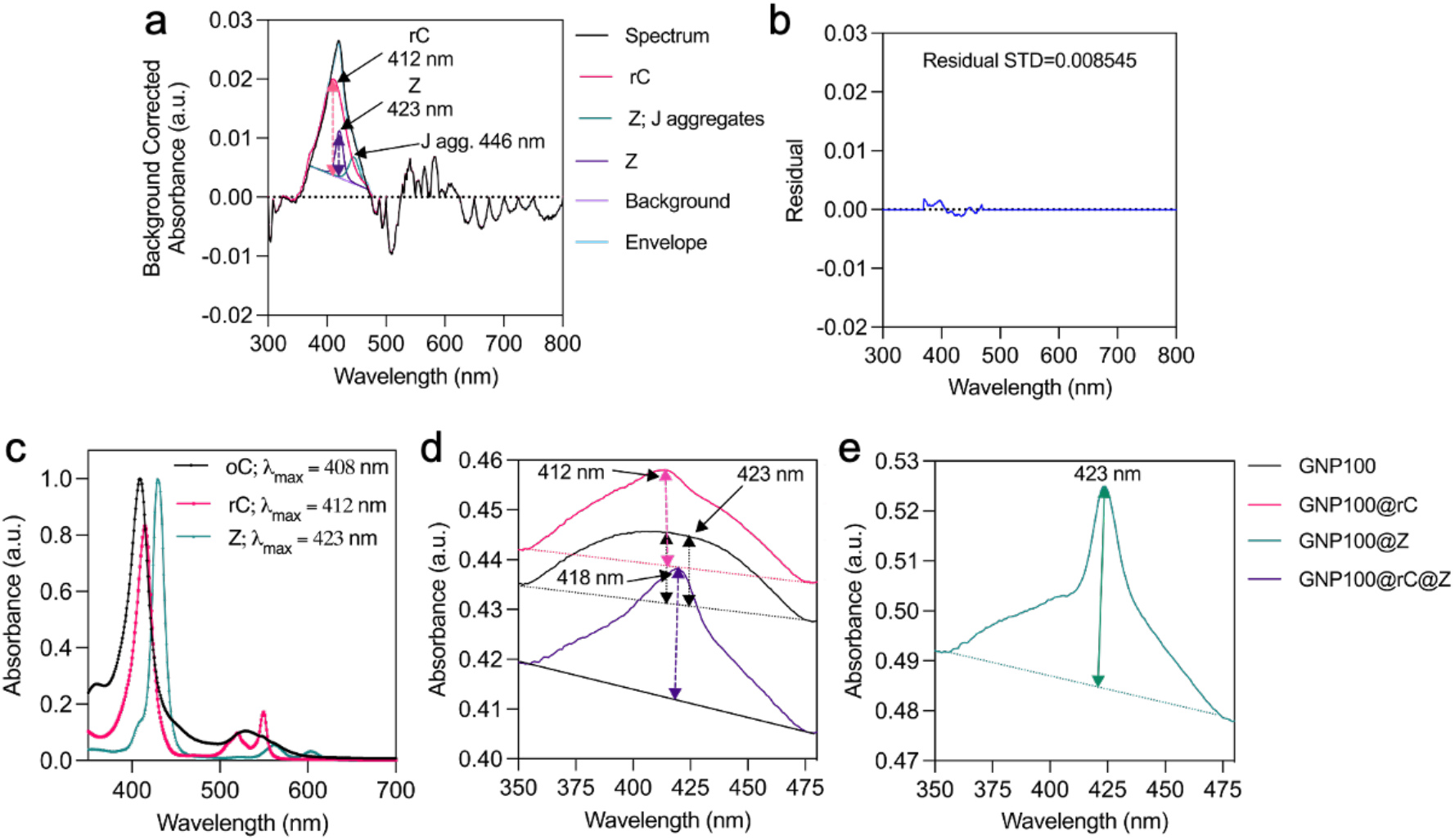
UV-vis absorption spectrum of bio-nanoantennae before electrical stimulation. **(a)** Deconvolution and fitting of UV-Vis spectrum of GNP100@rC@Z to identify and quantitate number of bound rC and Z molecules. **(b)** Residual standard deviation obtained after curve fitting. **(c)** UV-Vis spectrum of zinc porphyrin (Z), native cyt c in oxidised (oC) and reduced (rC) form, in PBS. **(d-e)** High resolution UV-Vis spectrum of bio-nanoantennae and baseline correction to quantify number rC and Z molecules bound to each nanoparticle (Supplementary Table **1-4**). **Note:** For quantification of rC and Z on GNPs, firstly a baseline correction factor was calculated by subtracting the absorbance of GNP100 at wavelengths 412 nm (rC) and 423 nm (Z) to the factor obtained by subtracting actual absorbance with baseline absorbance. Finally, this baseline correction factor was subtracted to obtain the final absorbance of rC and Z in GNP100@rC, GNP100@Z, GNP100@rC@Z samples. Absorbance final (Abs^f^) was taken to calculate cyt c concentration because of the influence on the spectra of the cyt c due to the SPR peak of GNP.

**Figure 4.**
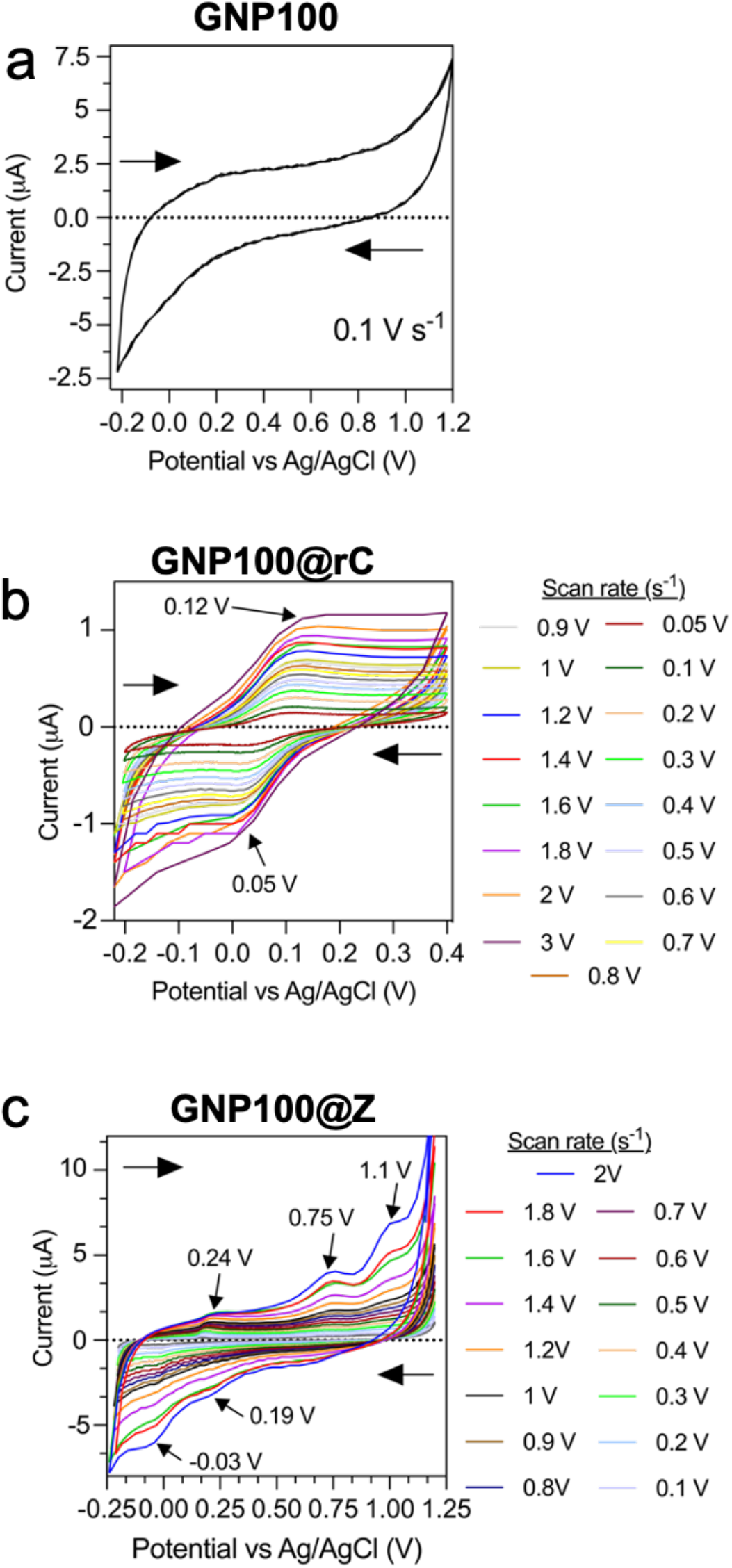
Cyclic voltammetry scan rate studies of **(a)** 100 nm GNPs capped with carboxylic-terminated PEG (GNP100), **(b)** GNP100 functionalised with reduced form of cyt *c* (rC) – GNP100@rC, and **(c)** Z functionalised GNP100 – GNP100@Z. Indium tin oxide (ITO) was used as a working electrode, counter electrode: platinum wire, and reference electrode: Ag/AgCl, electrolyte: 10 mM PBS. Sample concentration: 25 µg/mL.

**Figure 5.**
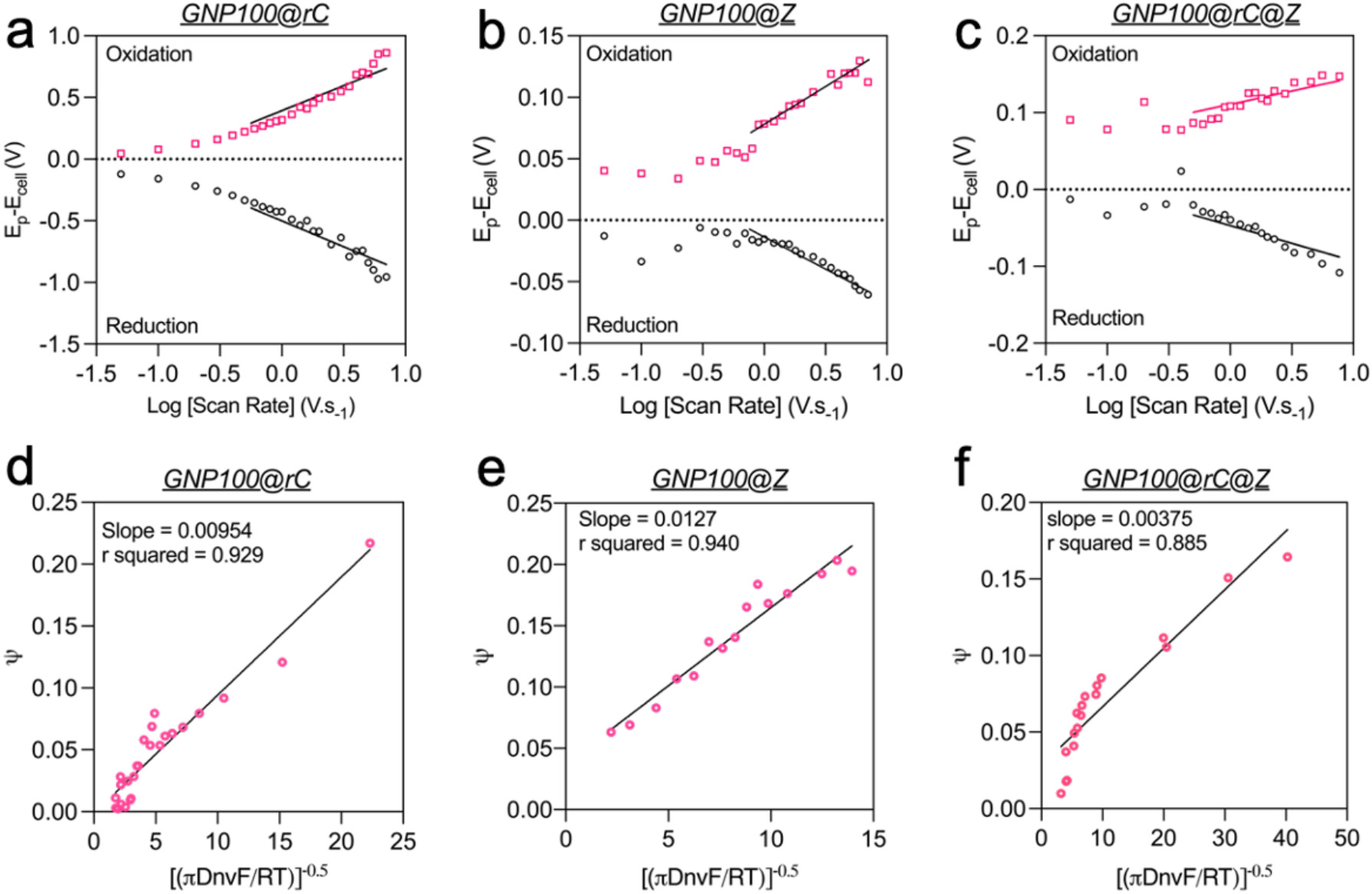
Determination of heterogenous rate constant (*k°)*. **(a-c)** Graph of peak potential – Standard Potential *vs* log of scan rate of different nanoantennae to determine rate transfer coefficient. **(d-f)** Nicholson plot to determine *k°* of nanoantennae. N = 3. Scan rates plotted are between 0.1 V/s and 3 V/s.

**Figure 6.**
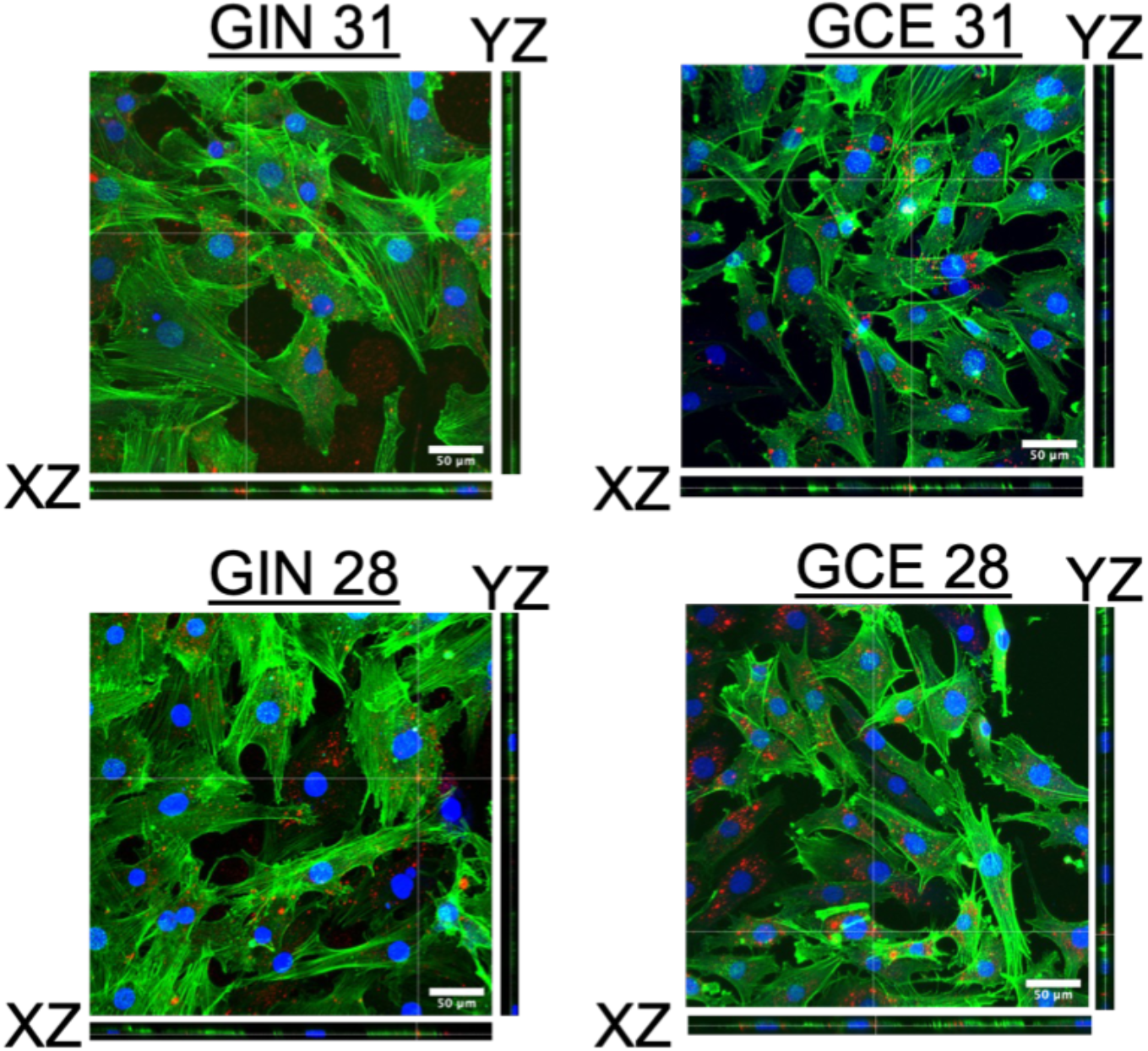
3D orthogonal Z-stack confirming the cytoplasmic localisation of bio-nanoantennae in GCE/GIN 31 and GCE/GIN 28 cells analysed using confocal microscopy. Cells were treated with bio-nanoantennae (GNP100@rC@Z) for 8 h and then fixed with paraformaldehyde followed by counterstaining with Cytopainter Actin Phalloidin (Alexa 488 green) stain and Hoechst nuclear stain (blue).

**Figure 7.**
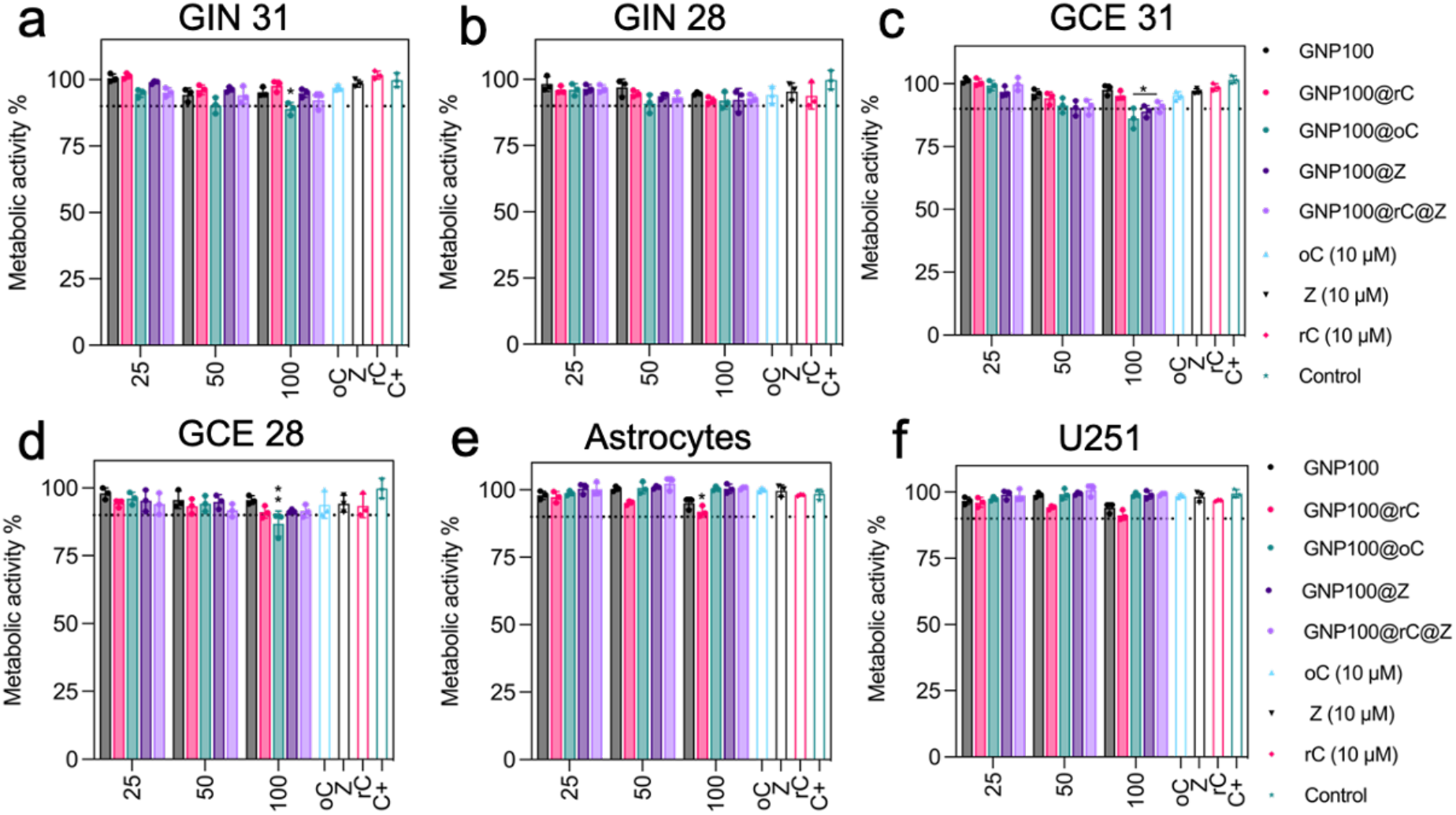
*In vitro* dose-dependent toxicity of different bio-nanoantennae synthesised using 2KDa PEG Linker. The cells were incubated with different nanocomposites at various concentration for 36 h before analysing their toxicity using PrestoBlue HS assay. **(a-f)** Metabolic activity of GIN 31, GIN 28, GCE 31, GCE 28, human derived cortical astrocytes, and U251 glioblastoma cells. GNP100 are 100 nm spherical gold nanoparticles functionalized with 2000 Da (2k) thiol-PEG-carboxylic, GNP100@rC were obtained by functionalising GNP100 with reduced form of cyt *c* (rC). Similarly, GNP100@oC, GNP100@Z, and GNP100@rC@Z are functionalised GNP100 with oxidised cyt *c*, zinc porphyrin (Z), and rC & Z, respectively. Error bars represent mean ± standard error of mean (S.E.M.) obtained from triplicate experiments repeated thrice. Statistical analysis was performed by applying 2-way ANOVA with a Tukey’s post-test.

**Figure 8.**
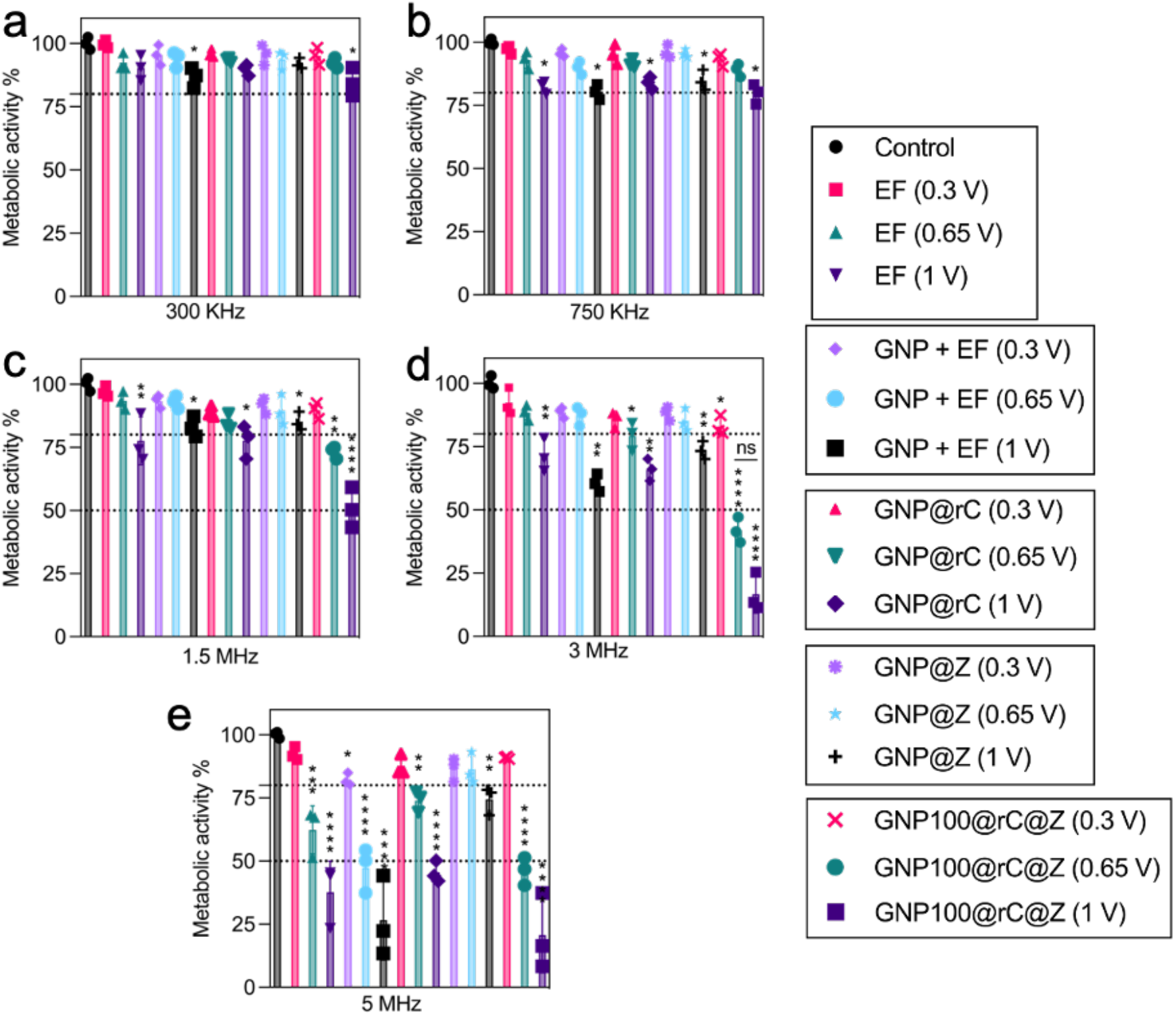
Electrical stimulation (ES) of GIN 31 cells to determine optimum AC-EF frequency and applied potential for bipolar electrochemistry mediated redox switching and activation of cyt *c* on the surface of bifunctionalised bio-nanoantennae. Metabolic activity of GIN 31 cells as a function of AC-EFs calculated using PrestoBlue HS assay. Cells were treated with nanoparticles for 8 h followed by AC-EFs stimulation of different frequency and voltages for 12 h. **(a)** 300 KHz, **(b)** 750 KHz, **(c)** 1.5 MHz, **(d)** 3 MHz, and **(e)** 5 MHz.

**Figure 9.**
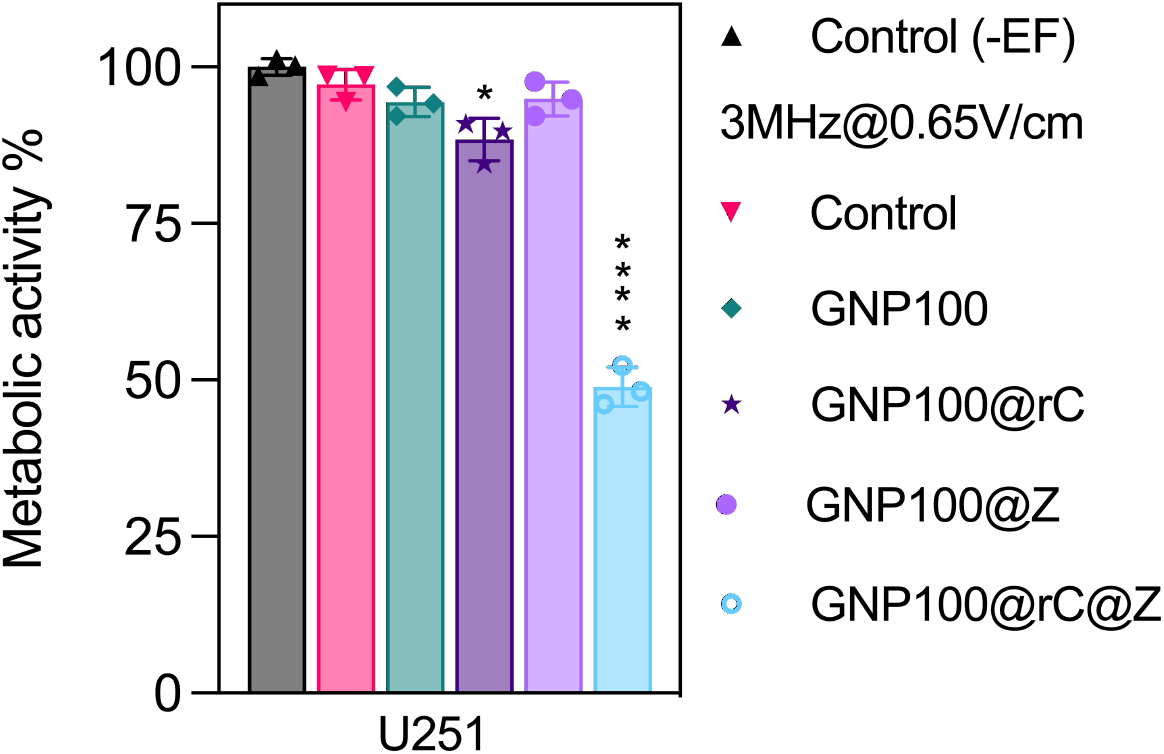
Bifunctionalised AC-EFs responsive bio-nanoantennae mediated wireless electrical molecular communication alters the metabolic activity of U251 cells. The cells were treated with GNP100@rC@Z for 8 h followed by 12 h treatment with AC-EF (3MHz, 0.65V/cm). Error bars represent mean ± standard error of mean (S.E.M.) obtained from triplicate experiments repeated thrice. Statistical analysis was performed by applying 2-way ANOVA with a Tukey’s post-test.

**Figure 10.**
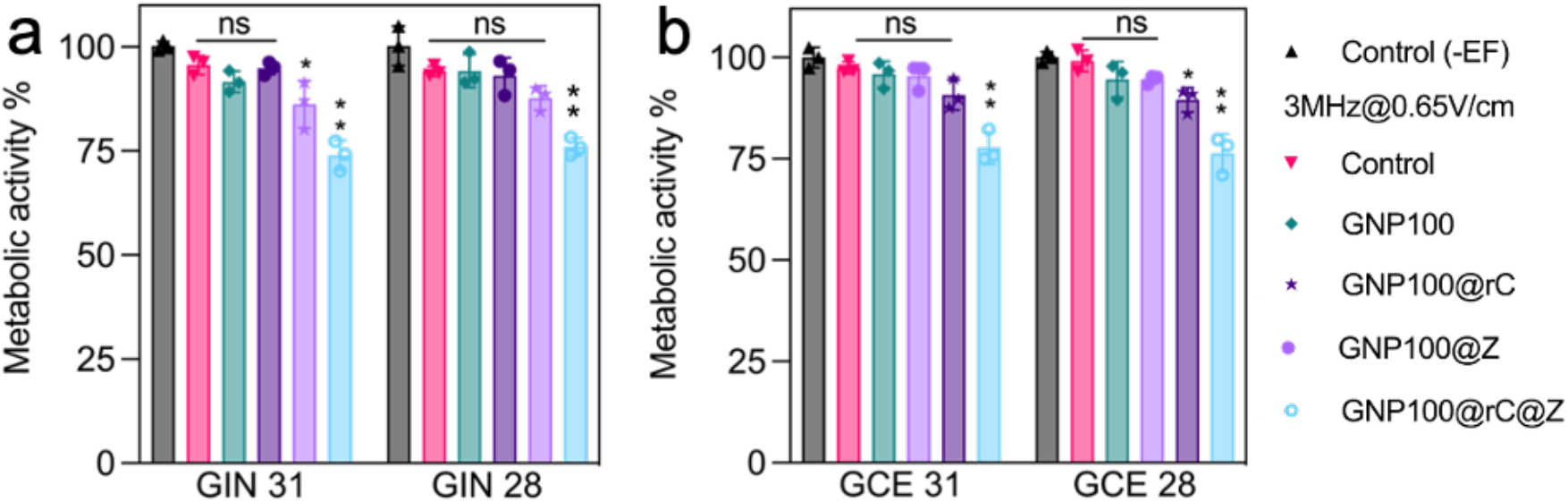
Bifunctionalised AC-EFs responsive bio-nanoantennae mediated wireless electrical molecular communication alters the metabolic activity of GBM cells. The cells were treated with GNP100@rC@Z for 8 h followed by 2 h treatment with AC-EF (3MHz, 0.65V/cm). Metabolic activity of **(a)** GIN, **(b)** GCE cells. Error bars represent mean ± standard error of mean (S.E.M.) obtained from triplicate experiments repeated thrice. Statistical analysis was performed by applying 2-way ANOVA with a Tukey’s post-test.

**Figure 11.**
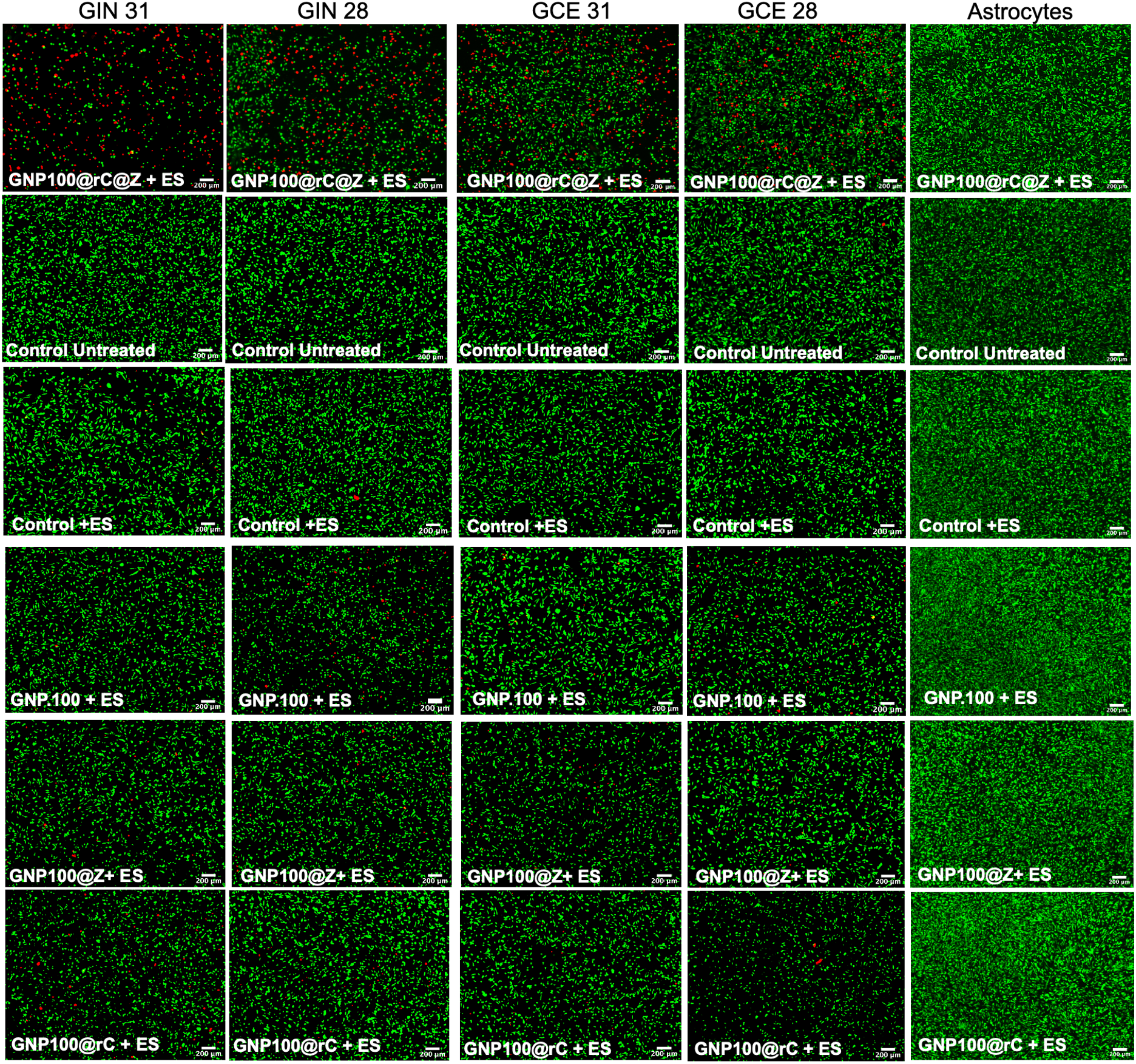
Live/dead imaging of GIN, GCE, and cortical astrocytes treated with bio-nanoantennae, and other control samples followed by AC EFs treatment for 12-hours. Post AC-EF treatment the cells were stained with calcein AM (green, live cells) and propidium iodide (red, dead cells). Live and dead cells were imaged using GFP and mTomato channel using a Nikon fluorescent microscope. Scale bar = 200 μm.

**Figure 12.**
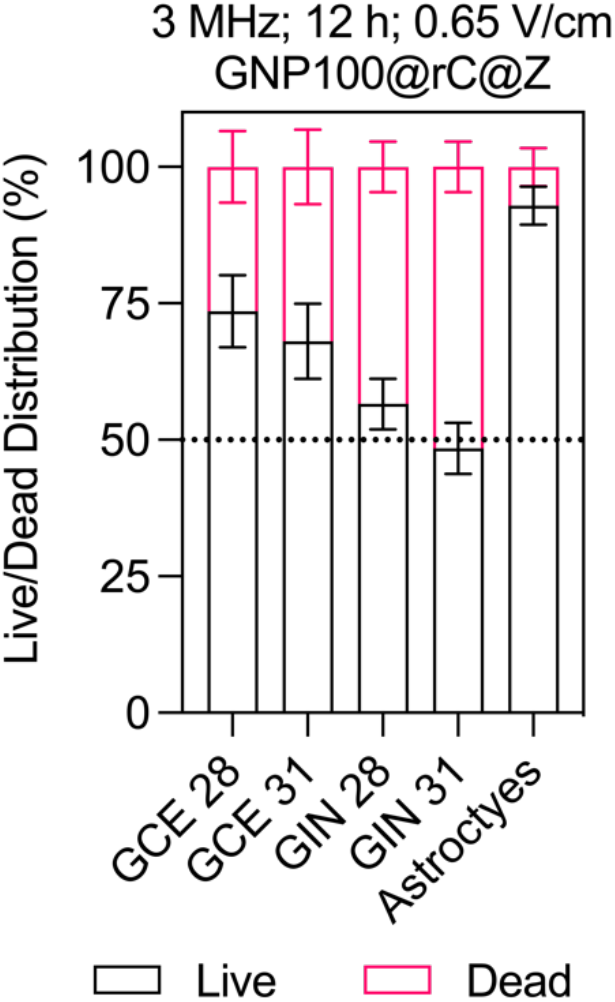
Quantification of live and dead cell population of images shown in supplementary fig. 11, calculated using ImageJ.

**Figure 13.**
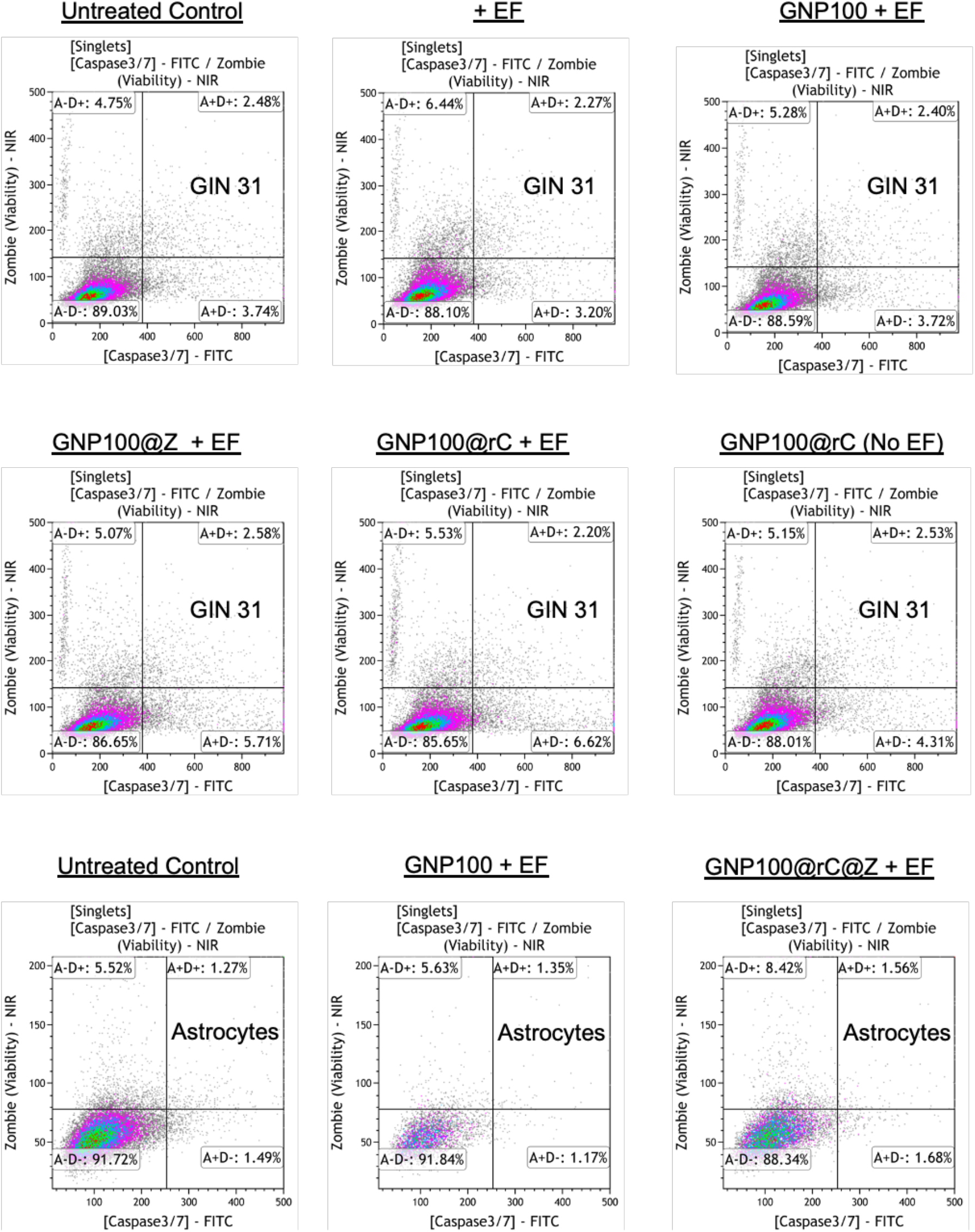
Representative flow cytometric analysis of GIN 31 cells (control samples) and cortical astrocytes (control and bio-nanoantennae) treated with CellEvent Caspase-3/7 Green to detect caspase activity and Zombie NIR dye to detect dead cell population after treatment with GNP100@rC@Z for 8 h followed by AC-EFs stimulation (3MHz, 0.65V/cm) for 12 h.

**Figure 14.**
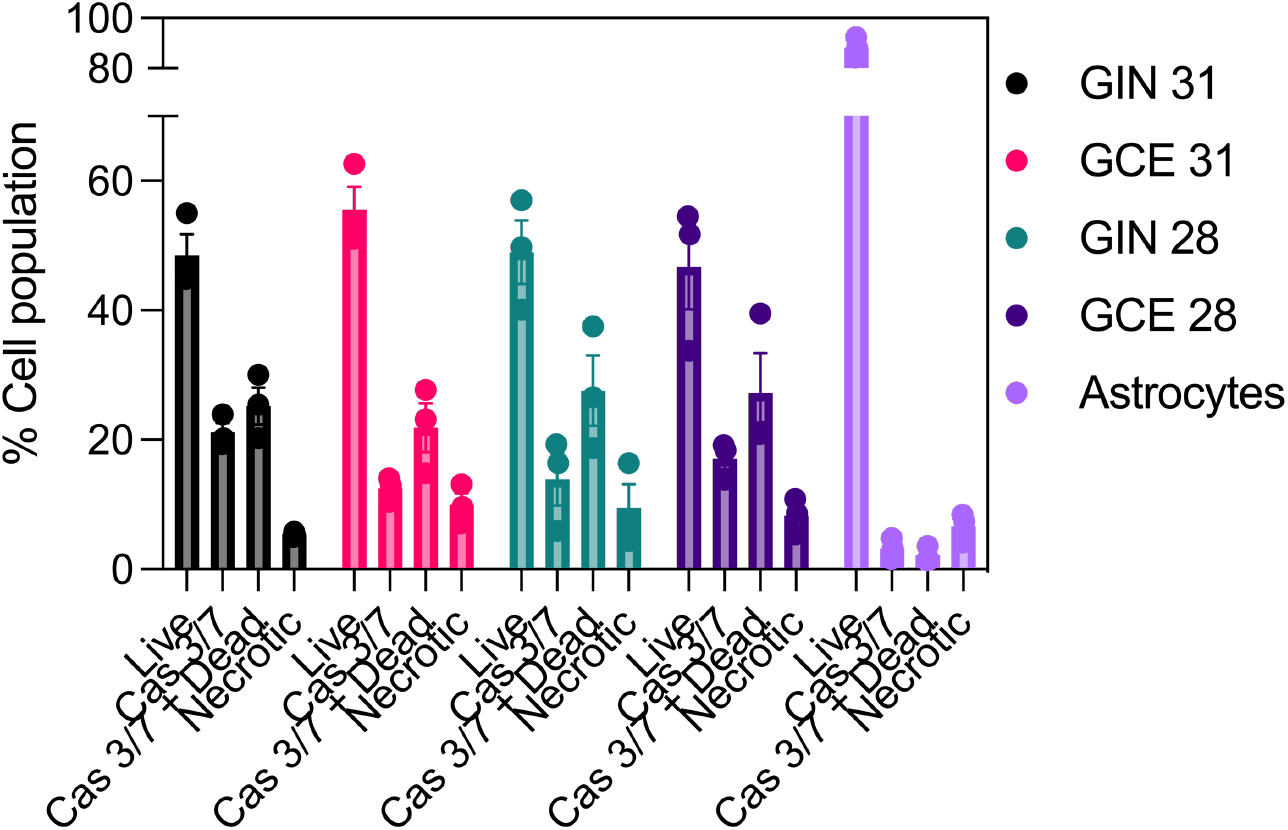
Quantification of flow cytometry data represented in Fig. 2d-g and supplementary Fig. 13. Error bars represent S.E.M. from triplicate experiments repeated thrice.

**Figure 15.**
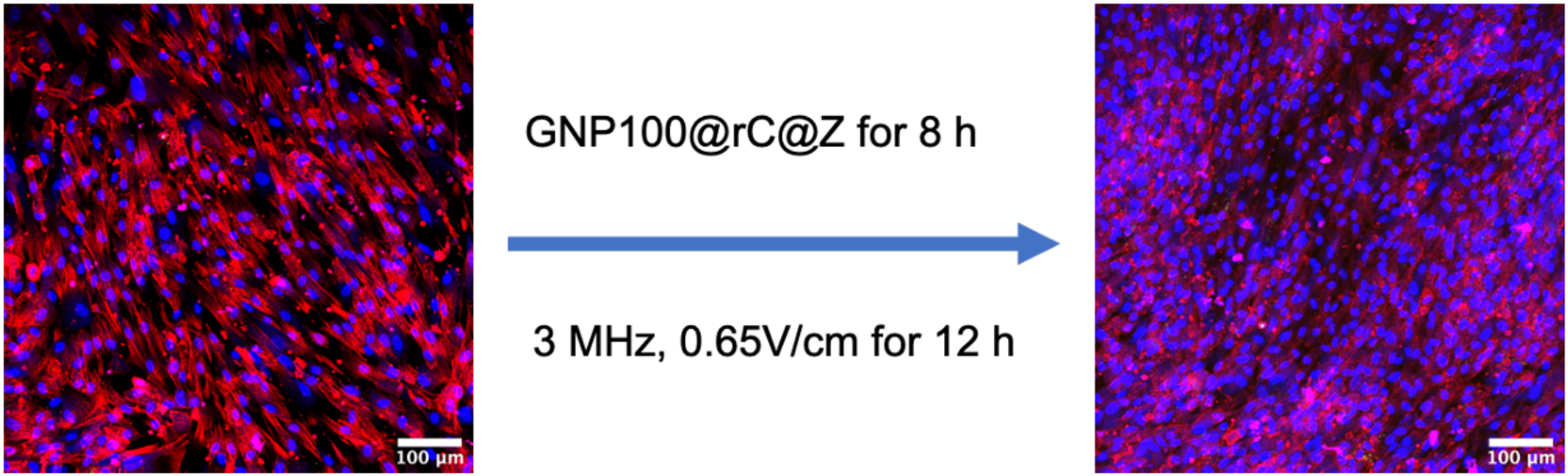
Confocal microscopy images to demonstrate caspase 3/7 activation in cortical astrocytes upon treatment bio-nanoantennae for 8 hours followed by stimulation with AC-EFs (3MHz, 0.65V/cm) for 12 h. Cells were fixed with paraformaldehyde followed by counterstaining with caspase 3/7 green detection kit, Cytopainter actin phalloidin (Texas Red 591 red), and Hoechst nuclear stain (blue). Scale bar = 100 μm.

**Figure 16.**
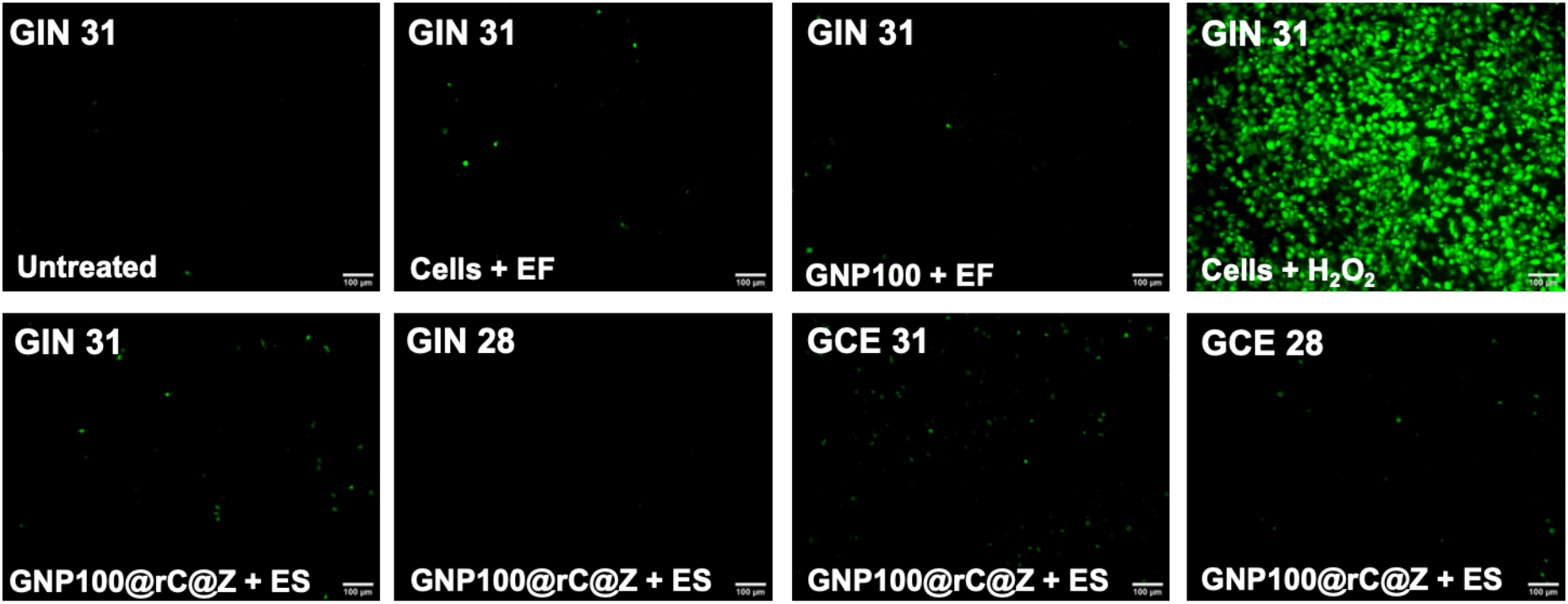
Representative fluorescent microscopy images of oxidative damage caused by bifunctionalised AC-EFs responsive bio-nanoantennae in GBM cells. GIN and GCE cells were treated with GNP100@rC@Z for 8 h followed by AC-EFs stimulation (3MHz, 0.65V/cm) for 12 h. Finally, the reactive oxygen generation was analysed using DCFDA/H_2_DCFDA - Cellular ROS Assay green detection Kit (ThermoFisher). Cells treated with 100 μM hydrogen peroxide for 30 minutes was used positive control. Scale bar = 100 μm.

**Figure 17.**
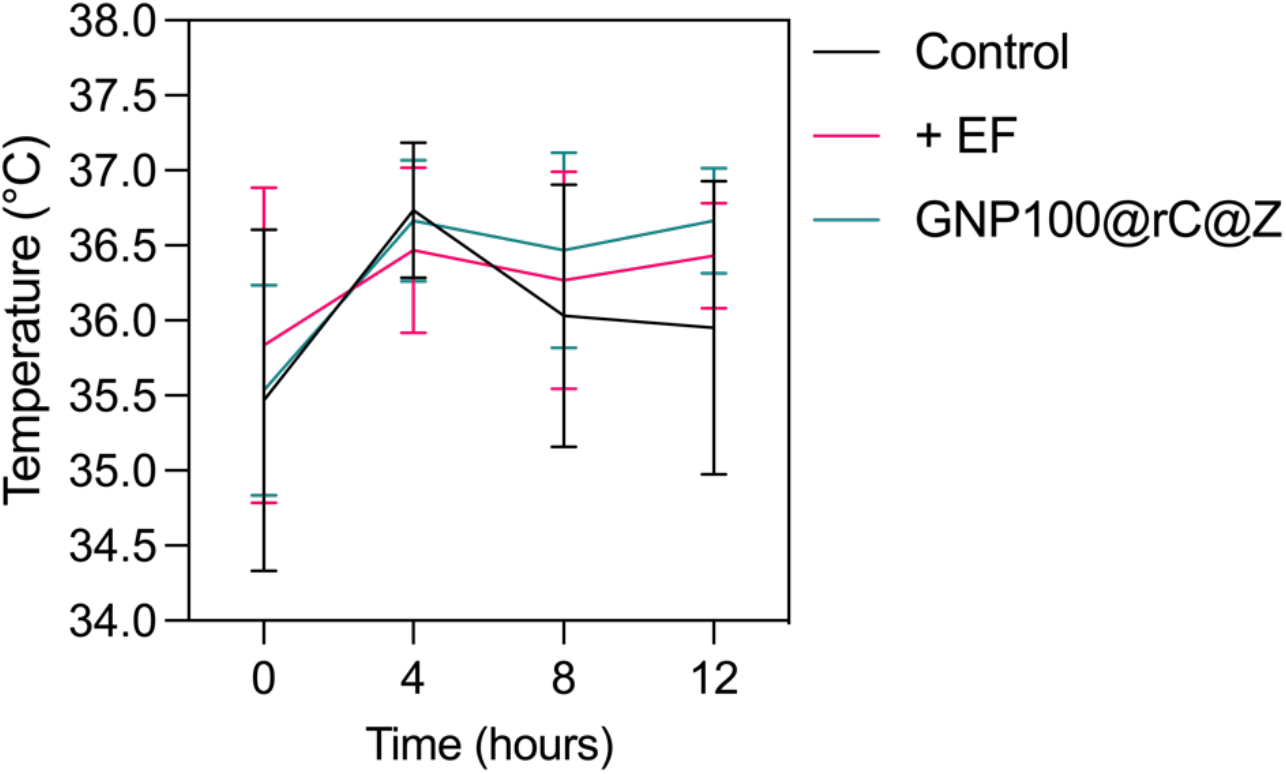
AC-EF (3MHz, 0.65V/cm) mediated change in solution temperature over the course of experiment measured using NIR laser gun.

**Figure 18.**
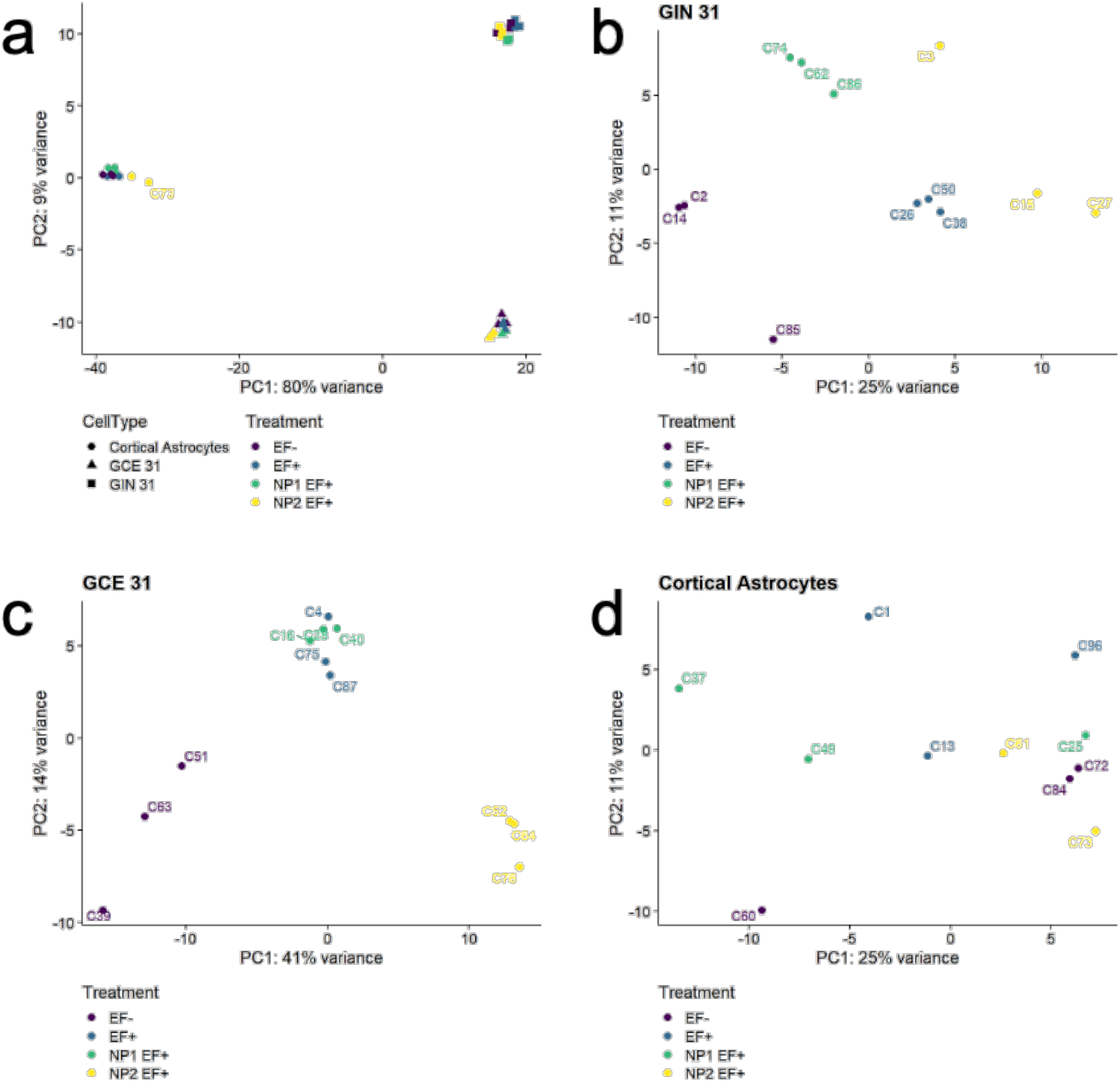
Principal component analysis (PCA). A variance stabilizing transformation was performed on the raw count matrix and 500 genes with the highest variance were used to plot the PCA. The variance was calculated agnostically to the pre-defined groups. PCA analysis **(a)** between different cell types and treatment groups, between different treatment within a cell type **(b)** GIN 31, **(c)** GCE 31, and **(d)** cortical astrocytes. In the figure treatment codes are as follow: - EF = Control (no treatment with either bio-nanoantennae or AC EFs); + EF = cells treated with AC EFs; NP1 EF+ = cell treated with GNP100@rC, and NP2 EF+ = cells treated with GNP100@rC@Z, for 8 h followed by 2-hour AC EFs.

**Figure 19.**
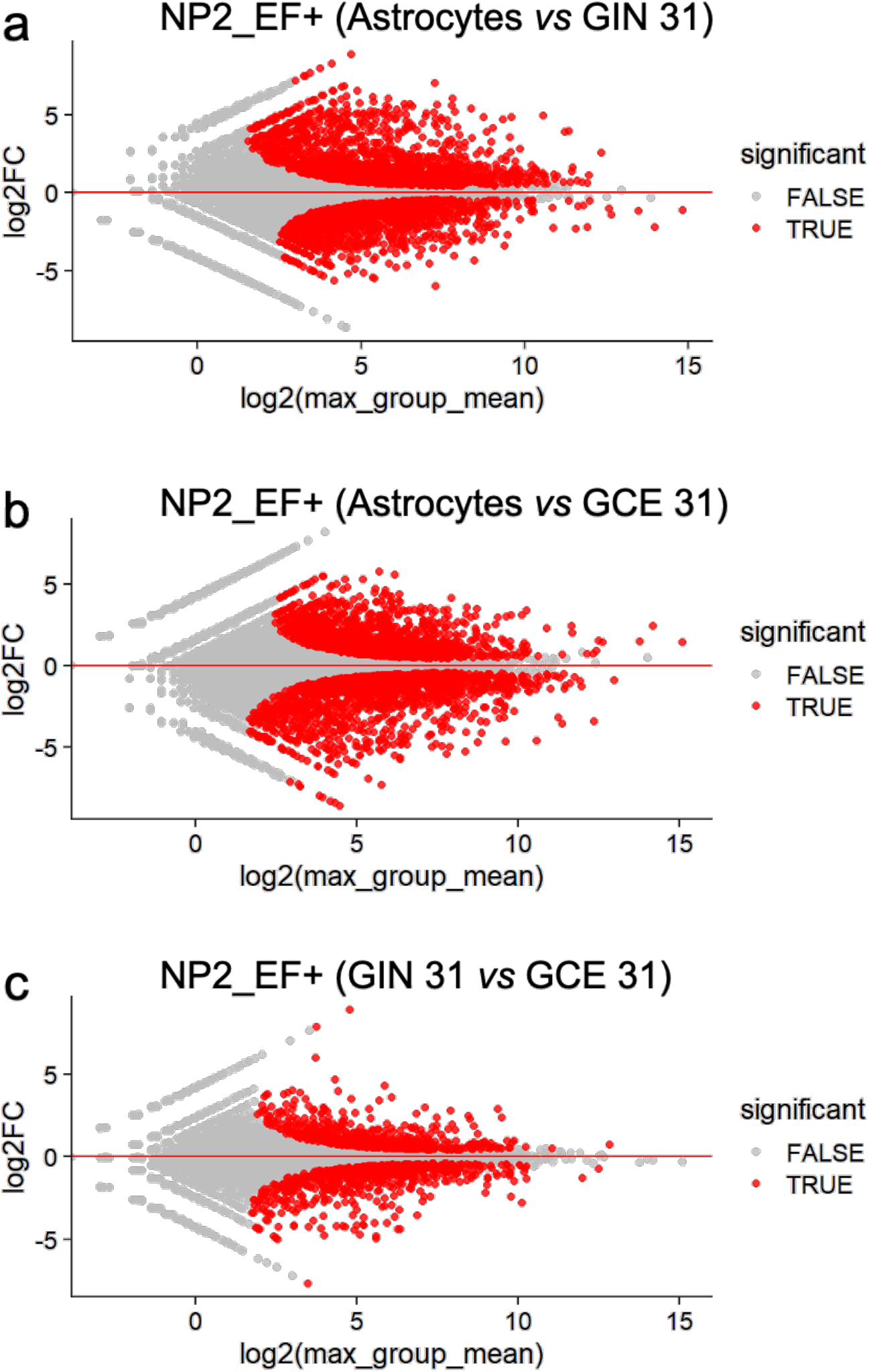
MA plot representing statistical tests (log fold-change *vs* mean expression between two cell types when treated with NP2_EF+) of the differential gene expression analyses. All significant differentially expressed genes are marked in red. (a) Cortical Astrocytes *vs* GIN 31, (b) Cortical Astrocytes *vs* GCE 31, and (c) GIN 31 *vs* GCE 31. Significant changes were defined as FDR<0.01. Note: NP2_EF+ = GNP100@rC@Z followed by ES with AC EFs 3MHz, 0.65V/cm for 2 hours.

**Figure 20.**
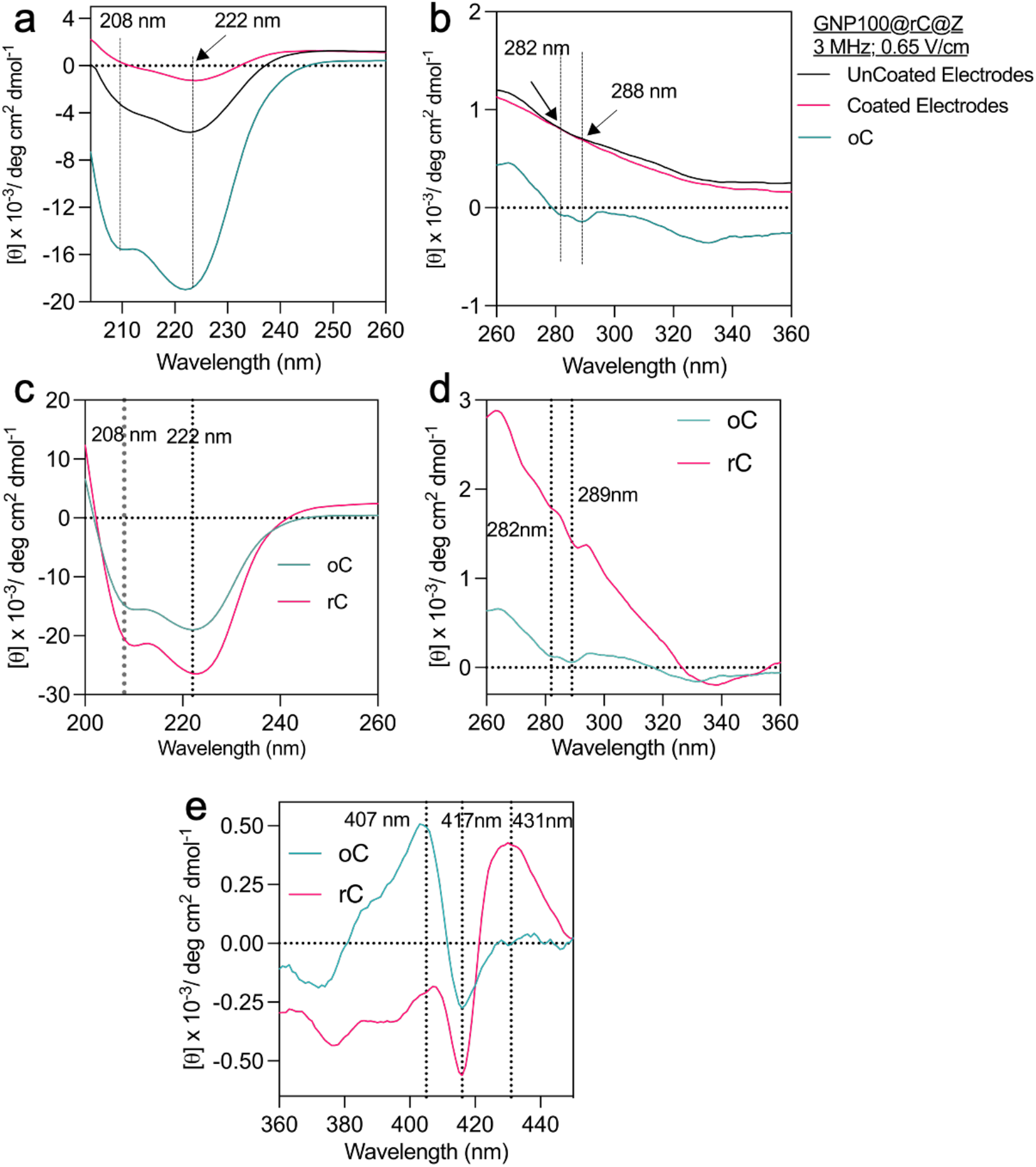
Circular Dichroism (CD) analysis. **(a-b)** Far-UV and near-UV CD of GNP100@rC@Z (bio-nanoantennae) upon exposure with AC EF_S_ of 3 MHz and 0.65 V/cm for 12 hours using coated and uncoated steel electrodes in PBS. The CD spectrum of bio-nanoantennae is compared with native oxidised cyt c (oC). **(c-e)** CD spectrum of native rC and oC in PBS with no ES.

**Figure 21.**
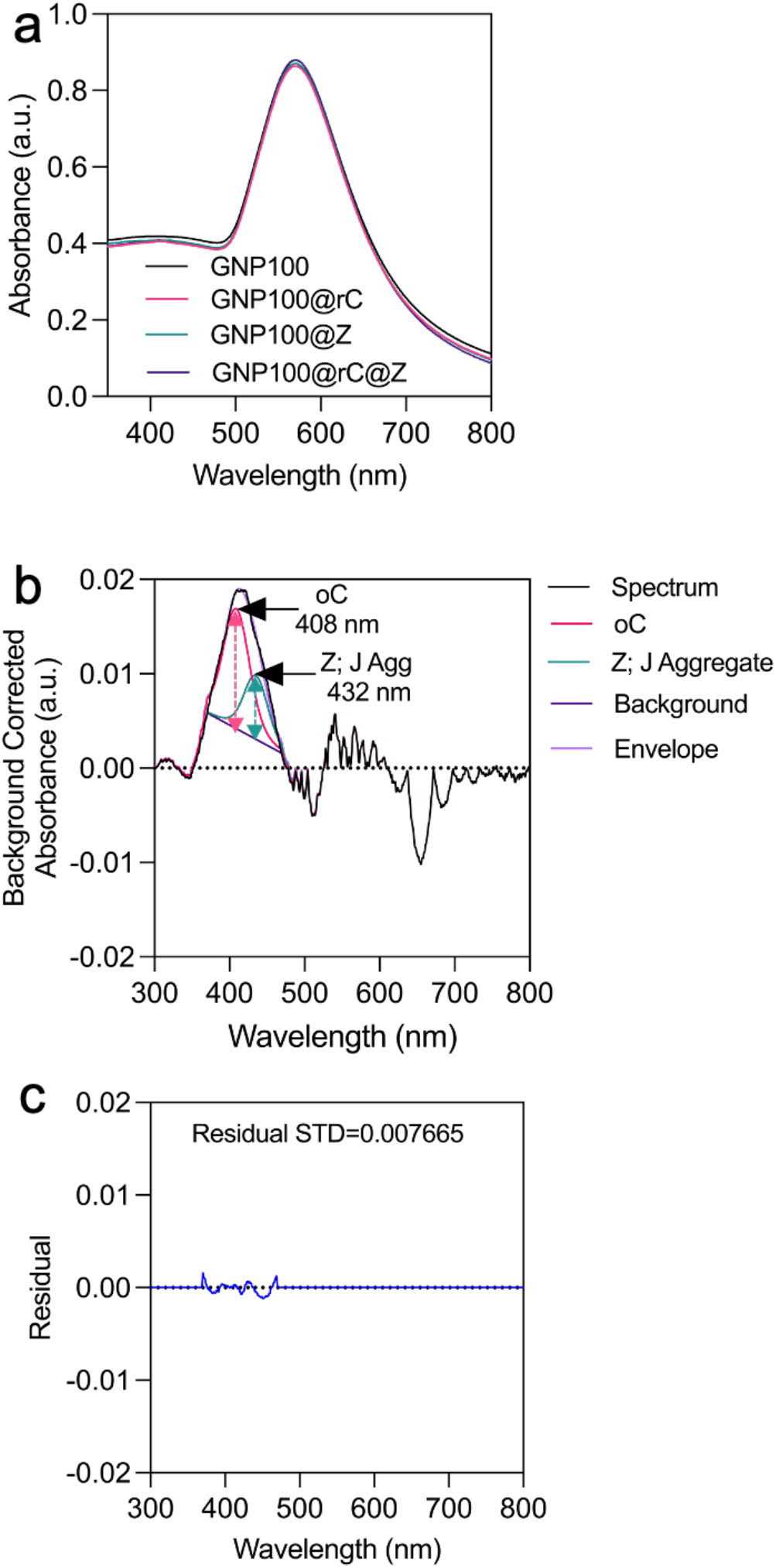
UV-Vis absorption spectrum of bio-nanoantennae after electrical stimulation with AC EFs (3 MHz, 0.65 V/cm). **(a)** Complete UV-Vis spectrum of different GNP conjugates of bio-nanoantennae. **(b)** High resolution deconvolution and curve fitting of GNP100@rC@Z samples confirming that electrical stimulation induces change in the redox state of rC to oC, and Z. **(c)** Residual standard deviation obtained after curve fitting

**Figure 22.**
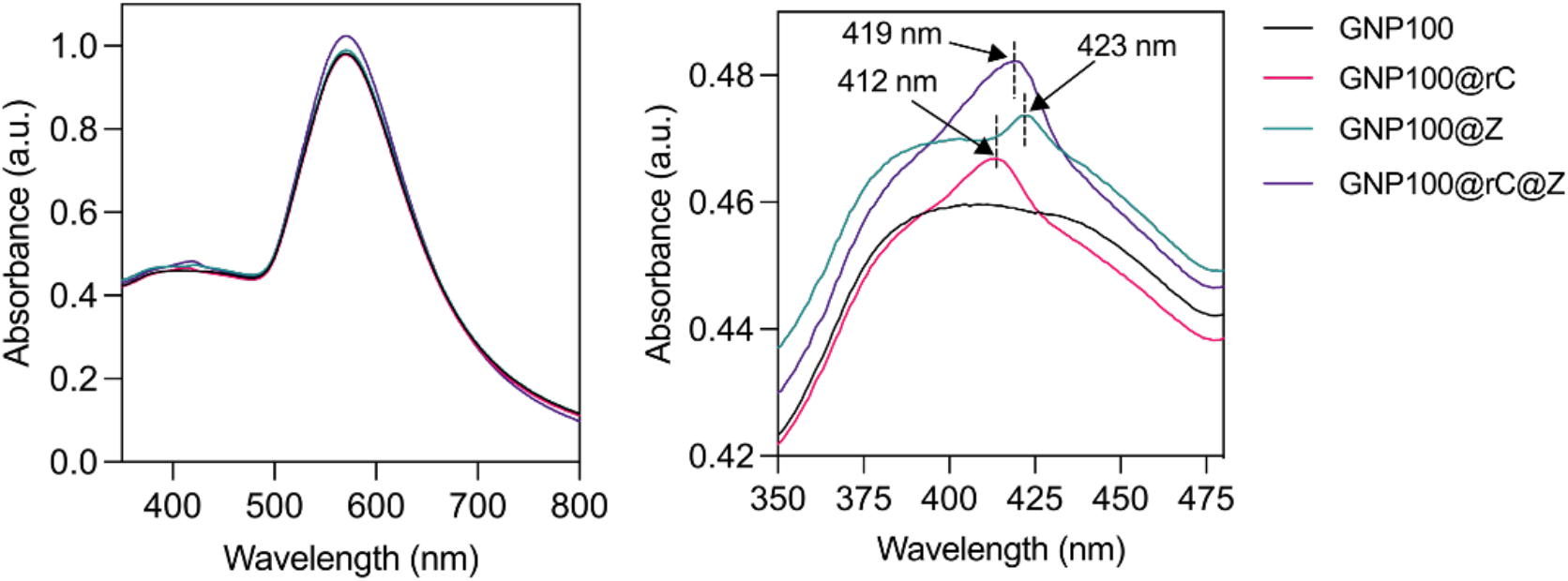
UV-Vis absorption spectrum of bio-nanoantennae without electrical stimulation after 12-hour incubation suggesting no change in redox state of rC and Z.

**Figure 23.**
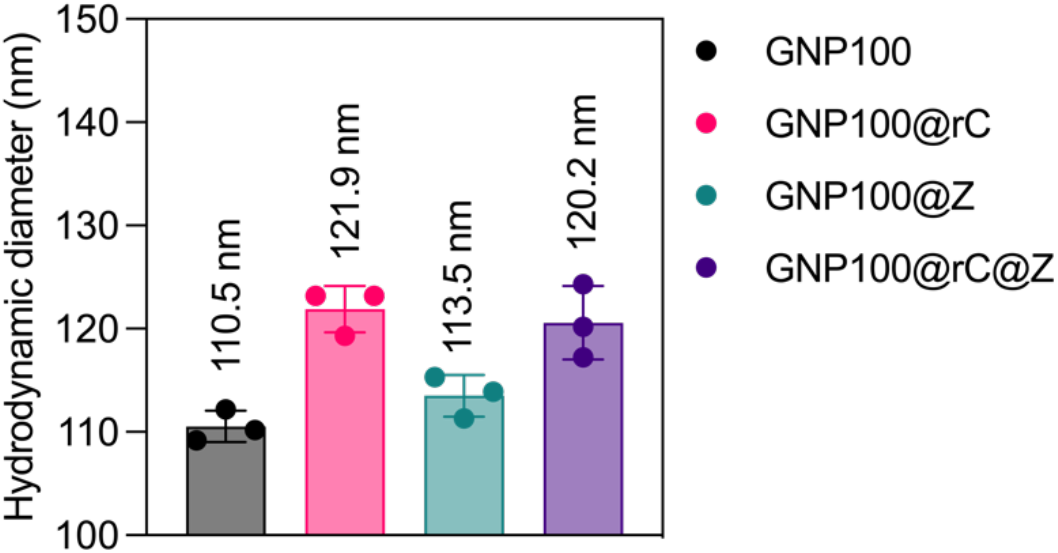
Hydrodynamic diameter of bio-nanoantennae (dispersed in ultra-pure water) after ES with AC-EFs (3 MHz, 0.65V). Error bars represent mean ± s.d. obtained from 3 individual experiments.

**Figure 24.**
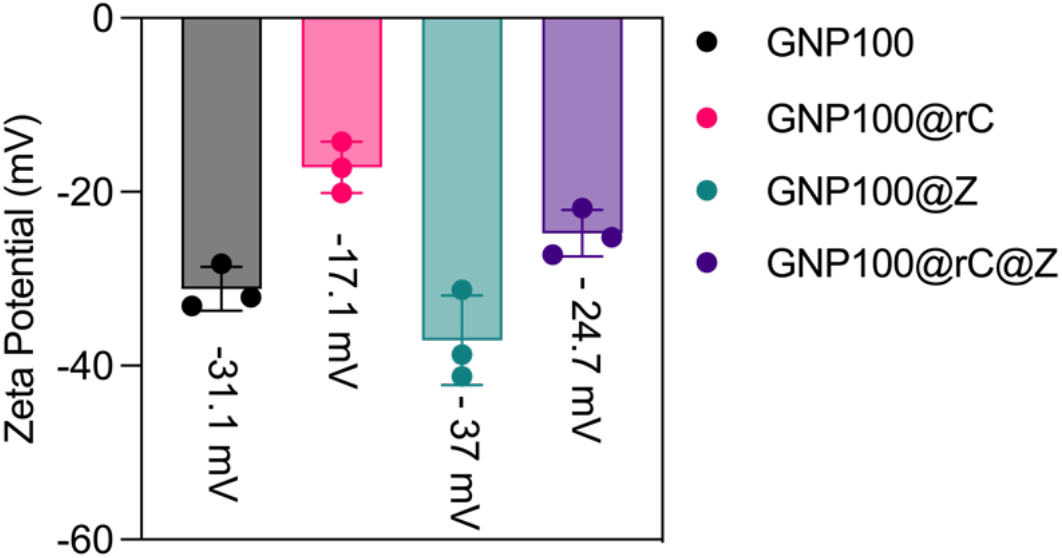
Zeta Potential of bio-nanoantennae (dispersed in ultra-pure water) after ES with AC-EFs (3 MHz, 0.65V). Error bars represent mean ± s.d. obtained from 3 individual experiments.

**Figure 25.**
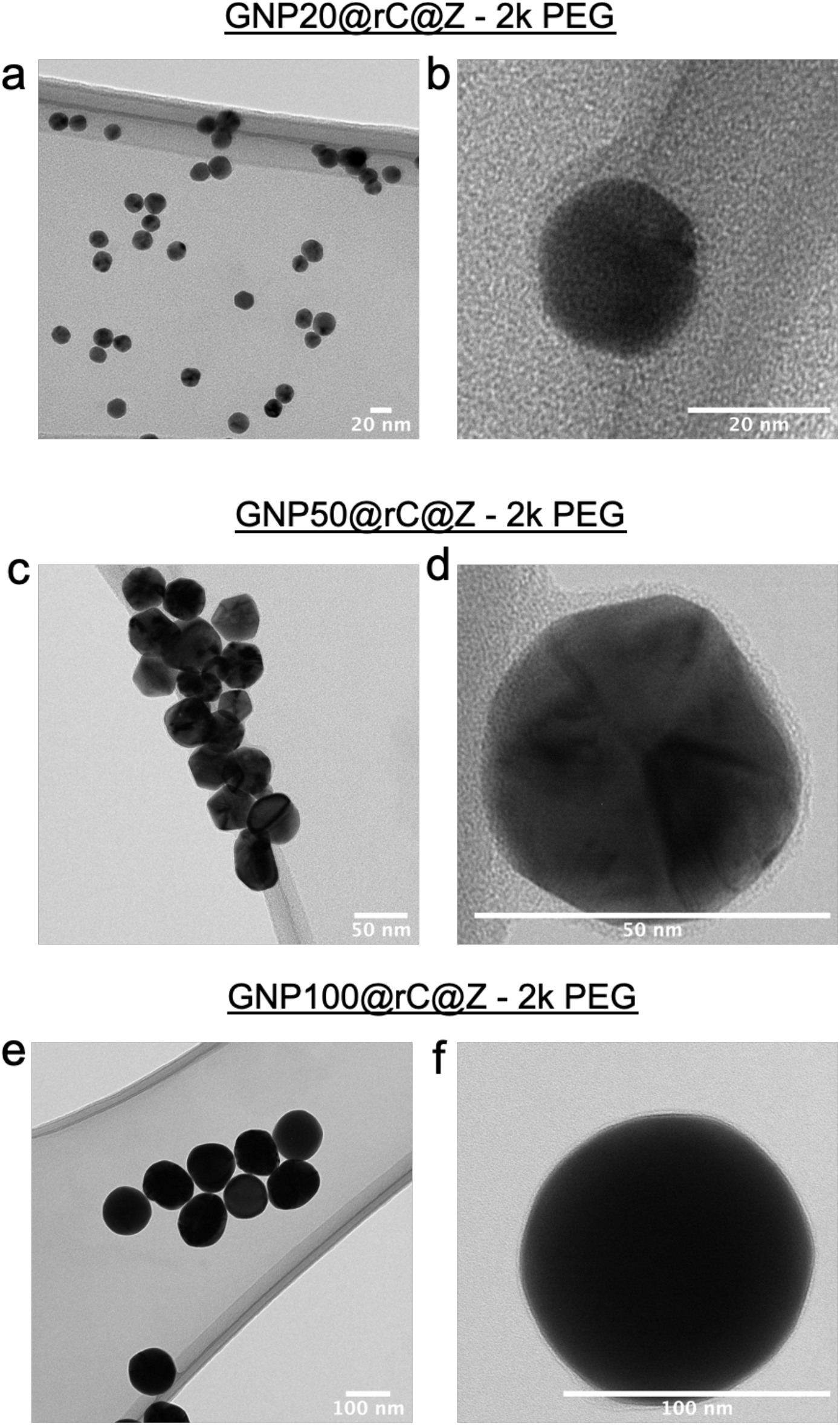
Transmission electron microscope images of different size bio-nanoantennae functionalised using 2000 Da thiol-PEG-carboxylic linker. **(a, c, and e)** TEM images of 20, 50, and 100 nm bifunctionalised bio-nanoantennae. **(b, d, and f)** High-resolution TEM images of 20, 50, and 100 nm bio-nanoantennae showing the presence of organic thiol-PEG linker.

**Figure 26.**
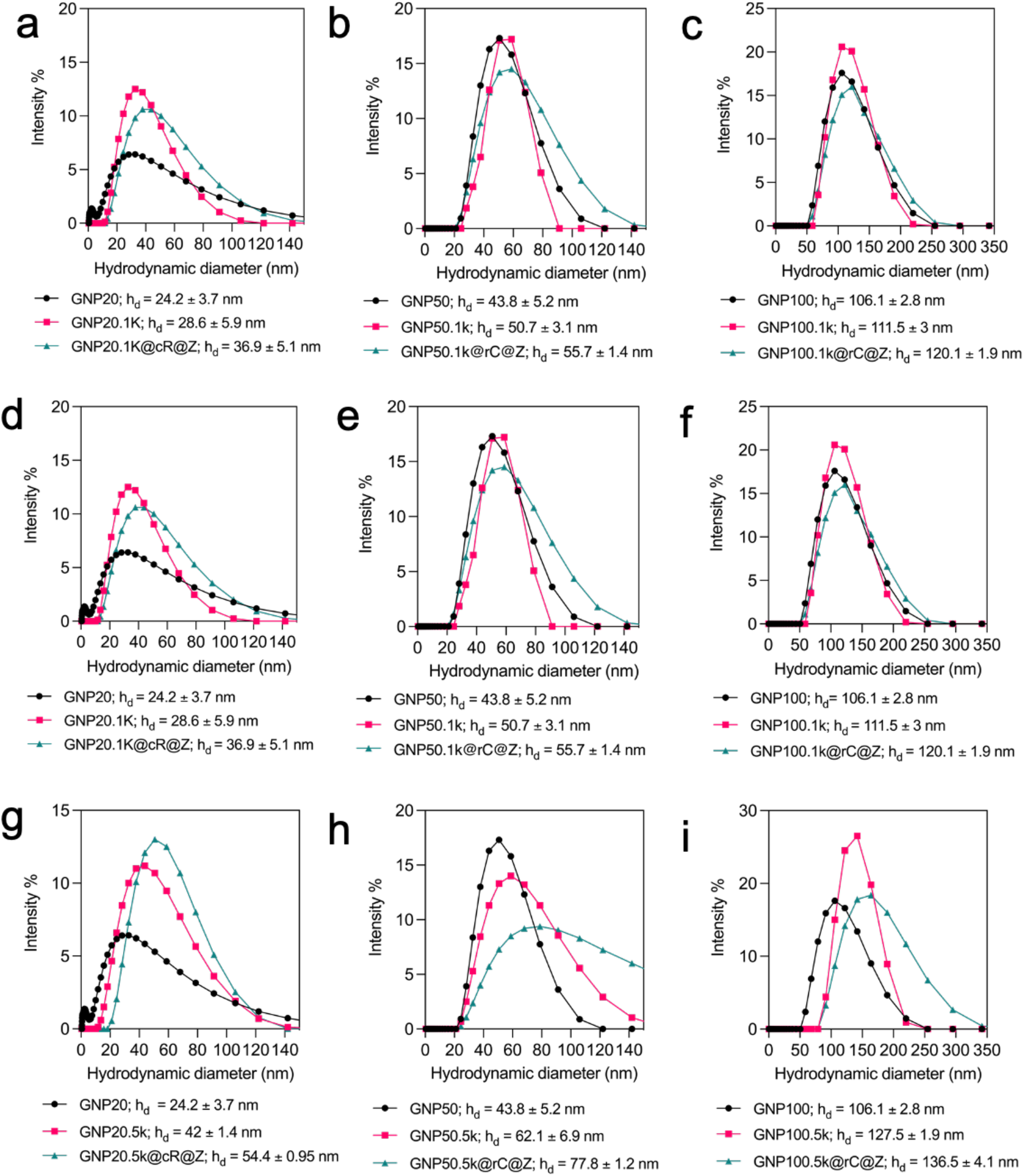
Size distribution of different size bio-nanoantennae (GNP20@rC@Z, GNP0@rC@Z, and GNP100@rC@Z) using linkers of various lengths analysed using dynamic light scattering (DLS). **(a-c)** Hydrodynamic diameter (h_d_) of bionanoatennae functionalised using 1000 Da (1k) **(d-f)** 2000 Da (2k) **(g-i)** Hydrodynamic diameter of 5000 Da (5k) thiol-PEG-carboxylic linker. Error bars represent mean ± standard error of mean (S.E.M.) obtained from triplicate experiments repeated thrice.

**Figure 27.**
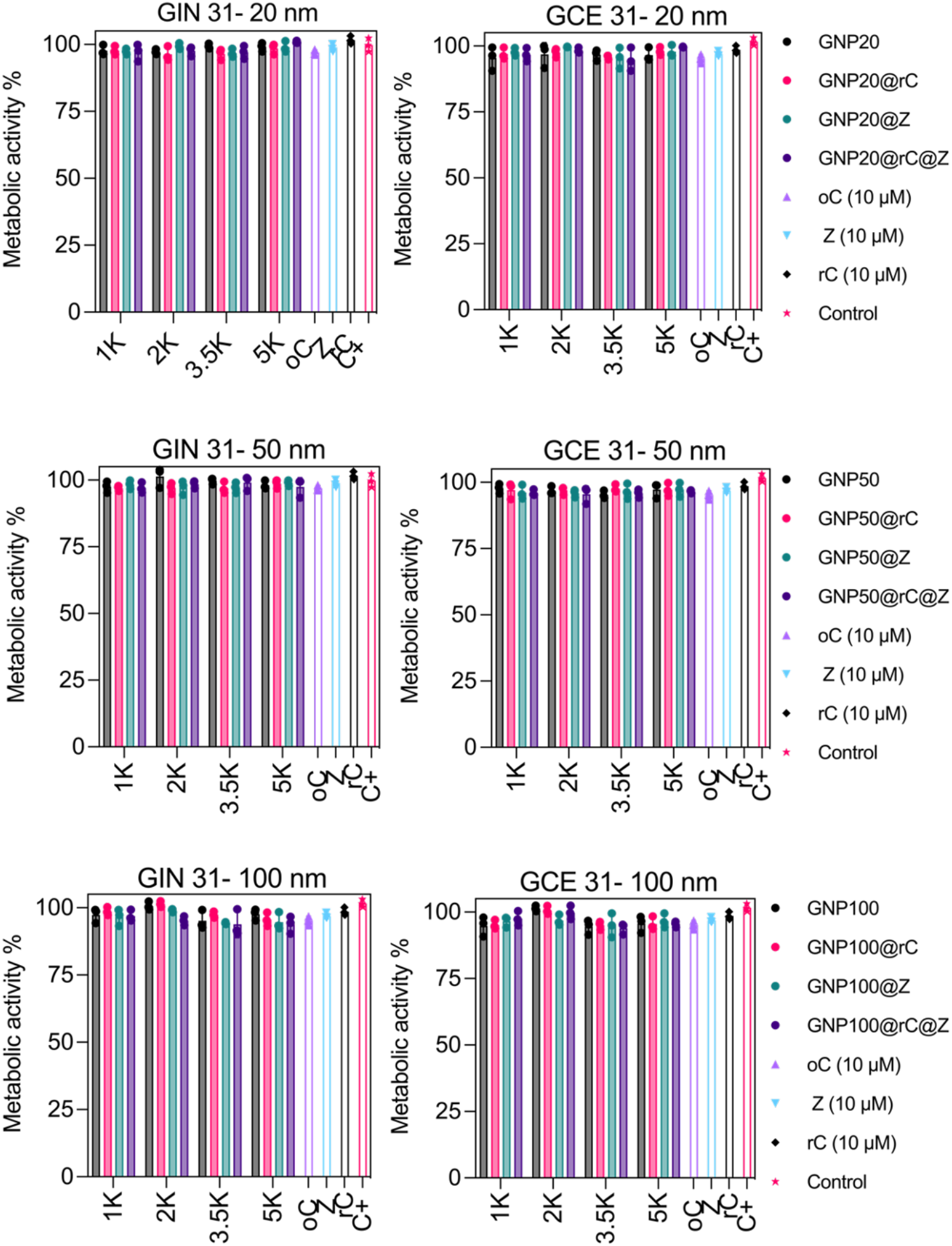
*In vitro* toxicity of GNP20, GNP50, and GNP100 bio-nanoantennae functionalised using PEG linker of different lengths in the absence of electric fields. The cells were incubated with different sized nanocomposites at a concentration of 25 μg/mL and their toxicity were analyzed using PrestoBlue HS cell viability kit. Metabolic activity of GIN 31 and GCE 31 upon treatment **(a-b)** 20 nm, **(c-d)** 50 nm, and **(e-f)** 100 nm with bio-nanoantennae functionalised using different thiol-PEG-carboxylic linker lengths. Error bars represent mean ± standard error of mean (S.E.M.) obtained from triplicate experiments repeated thrice. Statistical analysis was performed by applying 2-way ANOVA with a Tukey’s post-test.

**Figure 28.**
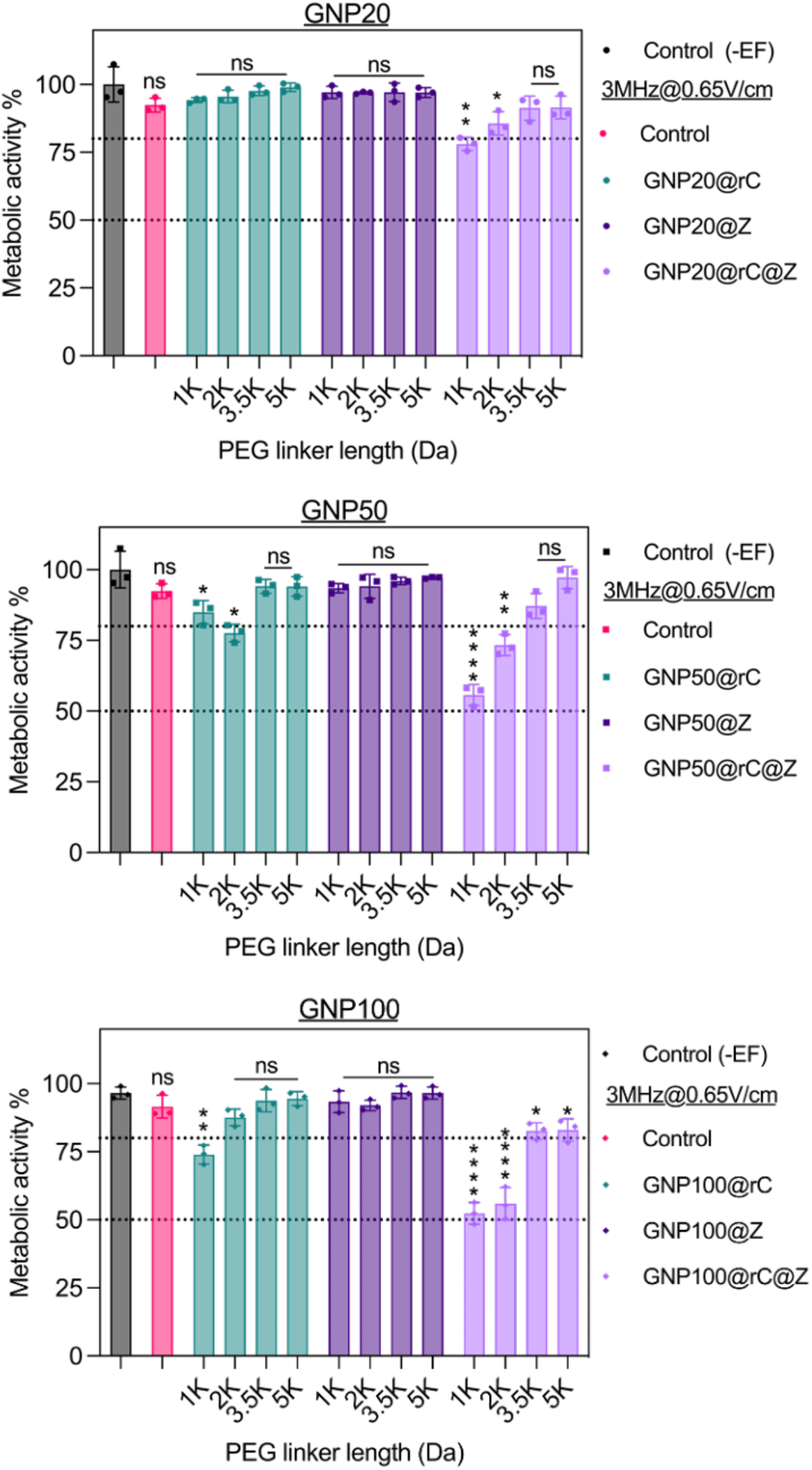
Wireless electrical-molecular quantum signalling: *in vitro* electron tunnelling via bio-nanoantennae for inducing cell death in GBM cells. Metabolic activity of GCE 31 cells as function of different sizebio-nanoantennae. (a) GNP20@rC@Z, (b) GNP50@rC@Z, and (c) GNP100@rC@Z synthesised using various linker lengths (1k, 2k, 3.5k, and 5k Da). GIN 31 cells were treated with bifunctionalised bio-nanoantennae for 8 h followed by AC-EFs stimulation (3 MHz, 0.65V/cm) for 12 h.

**Figure 29.**
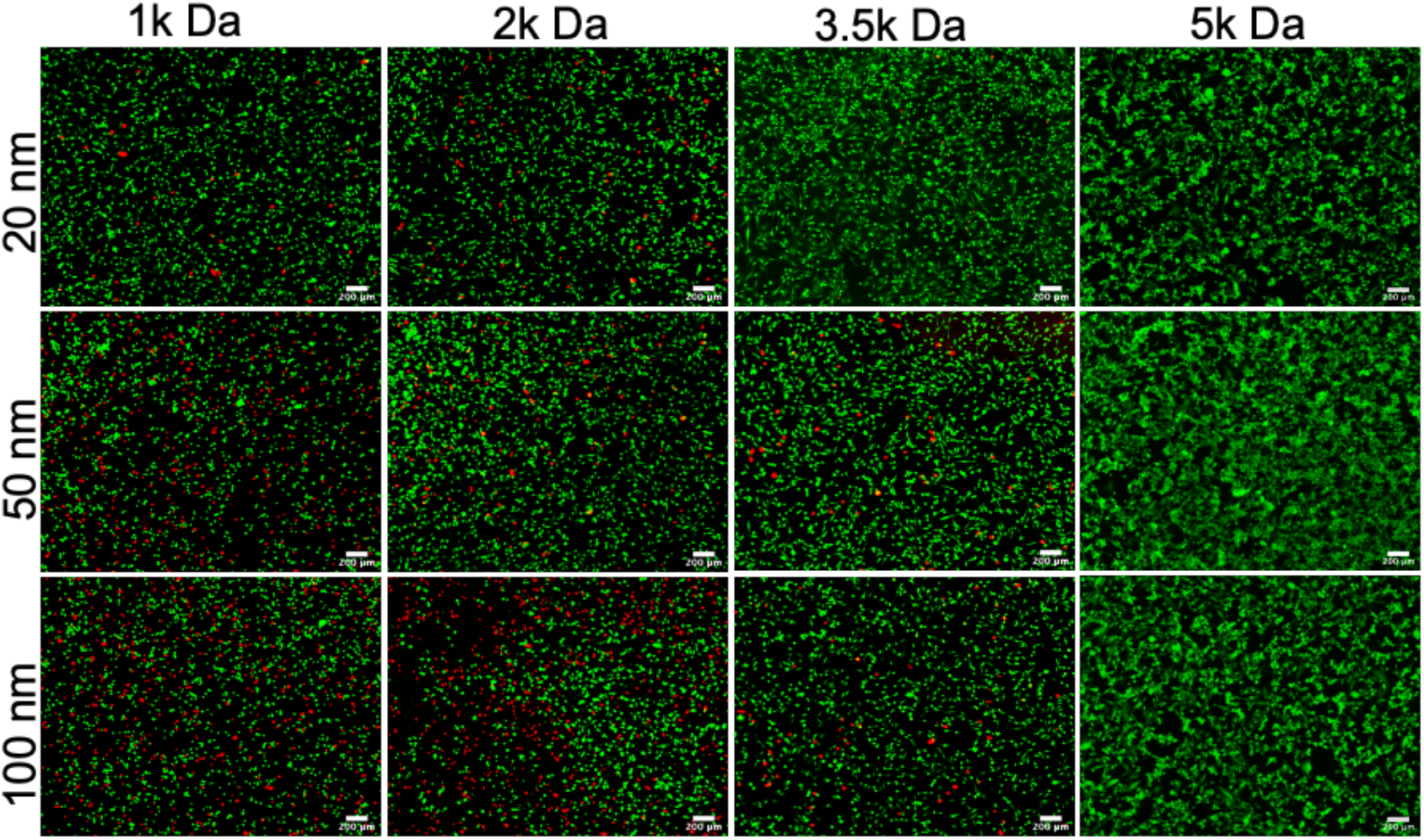
Live/dead staining to demonstrate the effect of linker length and bio-nanoantennae size on GIN 31 killing effect. Scale bar = 200 μm. GIN 31 cells were treated with bio-nanoantennae for 8 h followed by AC-EFs stimulation (3 MHz, 0.65V/cm) for 12 h. Post AC-EF treatment the cells were stained with calcein AM (green, live cells) and propidium iodide (red, dead cells). Live and dead cells were imaged using GFP and mTomato channel using a Nikon fluorescent microscope. Scale bar = 200 μm.

**Figure 30.**
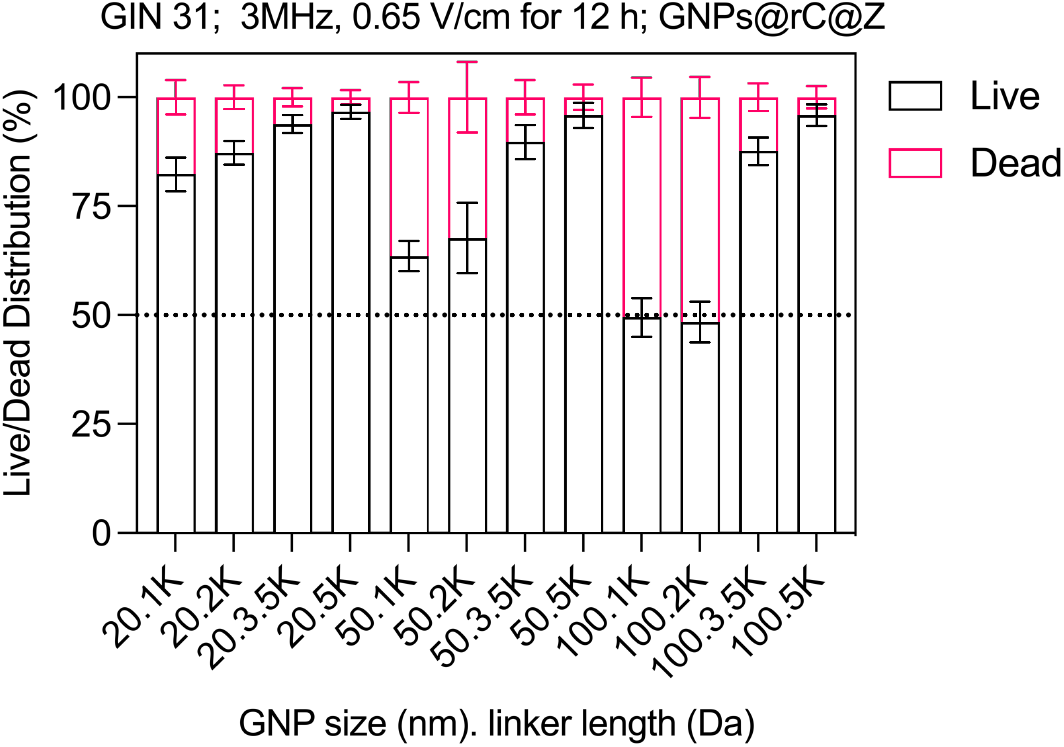
Quantification of live and dead cell population (shown in supplementary figure 29) to demonstrate the effect of linker length and bio-nanoantennae size on GIN 31 killing effect, calculated using ImageJ.

**Figure 31.**
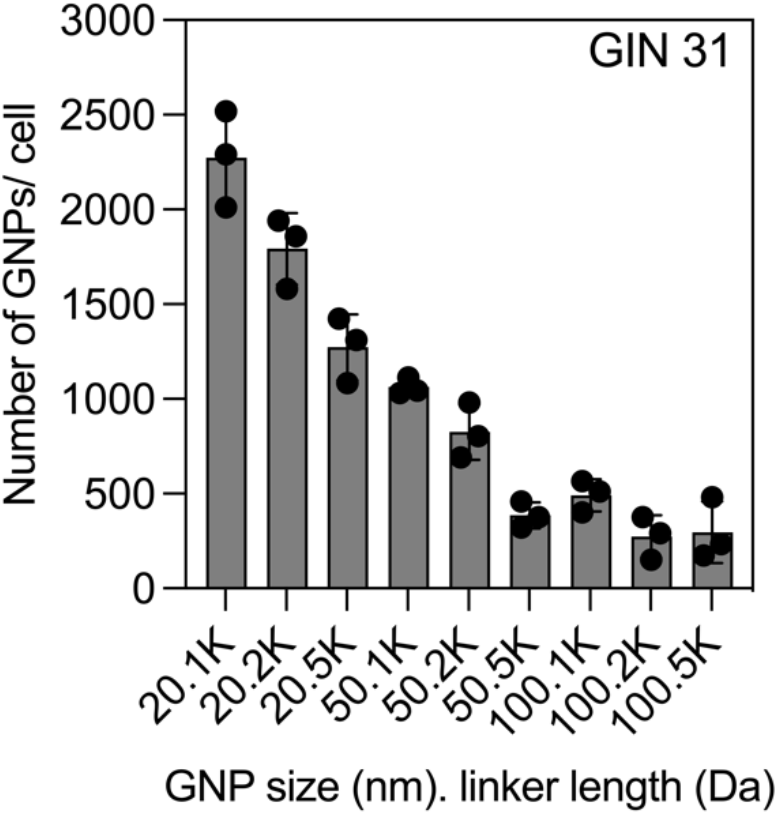
Number of bio-nanoantennae (different core and linker size) per GIN 31 cell calculated using ICP-MS to elucidate their cellular association.

**Figure 32.**
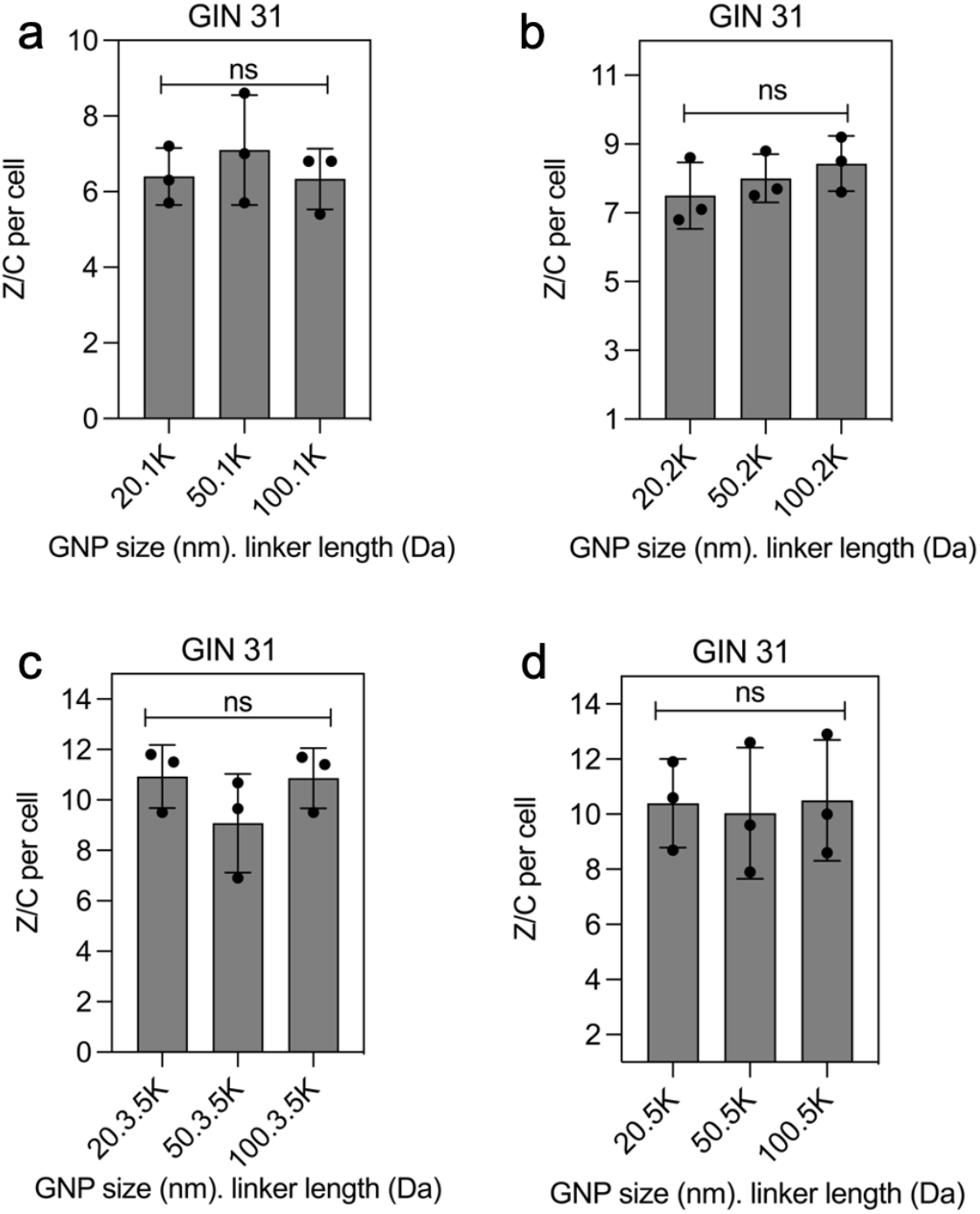
(a-d) Ratio of zinc porphyrin (Z) to cyt *c* (C) per GIN 31 cells determined from number of bio-nanoantennae per GIN 31 cells calculated using ICP-MS analysis.

**Figure 33.**
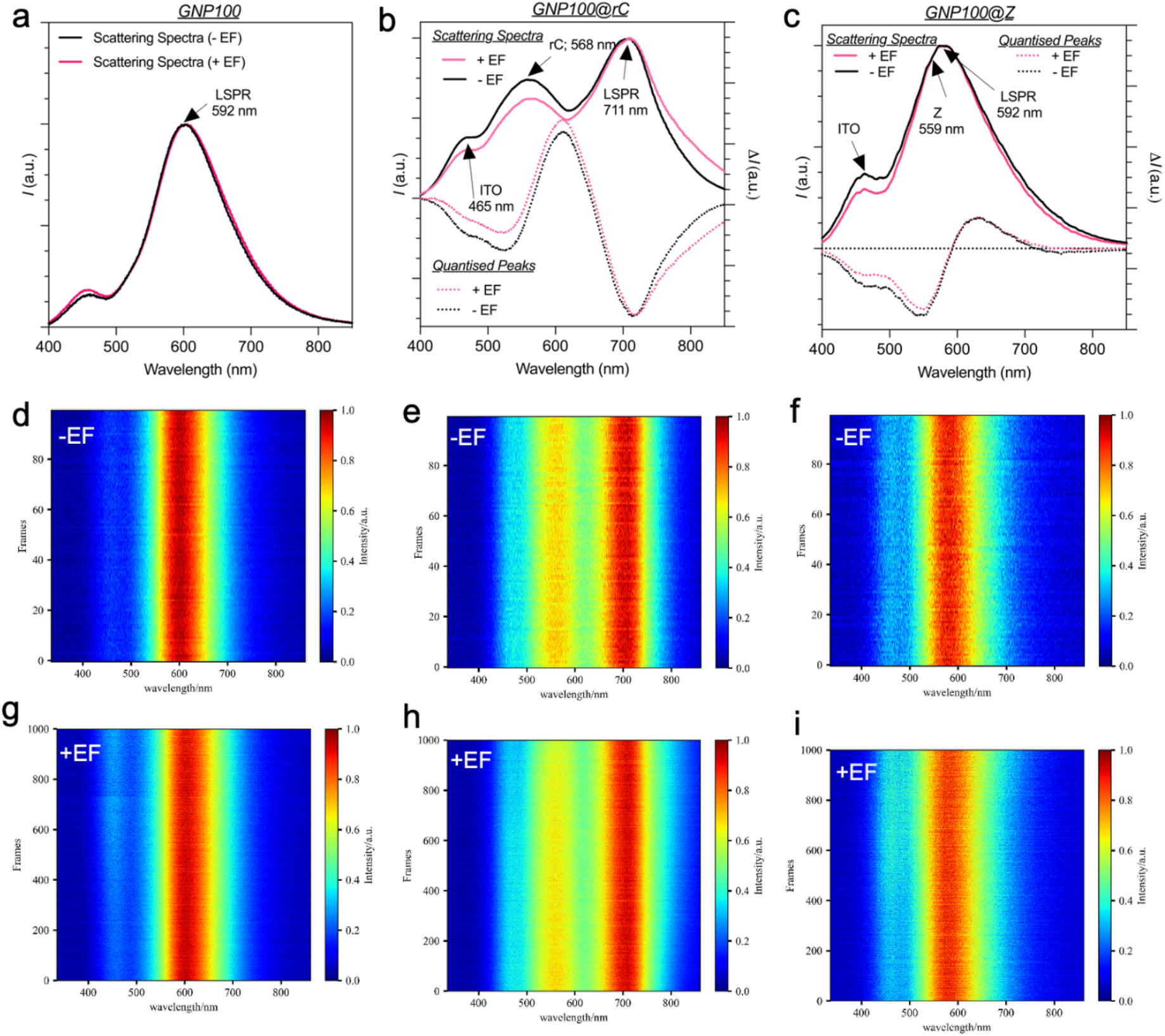
PRS scattering spectra and quantised peaks to demonstrated QBET. **(a-c)** Scattering spectra of GNP100 and corresponding 2D heat maps representing PRERT. **(d-f)** Scattering spectra and spectra difference for QBET obtained for GNP100@rC and **(g-i)** GNP100@Z. The quantised peaks were obtained from the difference of scattering spectra between the samples functionalised with rC, and Z using 2000 Da linker and GNP 100 nm. Solid curves are captured scattering spectra (linked to left axis), and dashed curves are quantised peaks i.e. the corresponding spectra difference (linked to right axis).

**Table 1.** Calculating the background correction factor (C.F.) from the UV Vis spectrum of GNP100@rC@Z before ES.

|  | B.C.rC |  | C.F.rC | B.C.Z |  | C.F.Z |
| --- | --- | --- | --- | --- | --- | --- |
| Sample | Abs $\lambda$<br>412 nm | Abs $\lambda$ =<br>412 nm<br>baseline | rC | Abs $\lambda$<br>423 nm | Abs $\lambda$ =<br>423 nm<br>baseline | Z |
| GNP100@rC@Z | 0.445 | 0.431 | 0.014 | 0.445 | 0.42 | 0.025 |

Where,
**B.C.rC** = Background correction for rC; **C.F.rC** = Correction factor for rC
**B.C.Z** = Background correction for Z; **C.F.Z** = Correction factor for Z

**Table 2.** Calculating the concentration of rC and Z bound to single Gold Nanoparticle before ES was calculating using Beer-Lamberts law.

| Sample | rC.Abs<sub>a</sub> | Z.Abs<sup>a</sup> | rC.Abs<sup>b</sup> | Z.Abs<sup>b</sup> | rC<br>Abs<sup>f</sup> | Z<br>Abs<sup>f</sup> | Concentration<br>(M) |
| --- | --- | --- | --- | --- | --- | --- | --- |
| GNP100@rC | 0.457 | N/A | 0.438 | N/A | 0.005 | N/A | $3 \times 10^{-8}$ |
| GNP100@Z | N/A | 0.523 | N/A | 0.487 | N/A | 0.011 | $1.9 \times 10^{-7}$ |
| GNP100@rC@Z | 0.019 | 0.01 | 0.003 | 0.003 | 0.002 | 0.007 | rC = $1.3 \times 10^{-8}$<br>Z = $1.2 \times 10^{-7}$ |

Where,
**rC.Abs<sup>a</sup>** and **Z.Abs<sup>a</sup>** = Actual absorbance of rC and Z at 412 and 423 nm in Supplementary Fig. 3d, respectively.
**rC.Abs<sup>b</sup>** and **Z.Abs<sup>b</sup>** = Absorbance at the baseline of rC and Z.
**rC.Abs<sup>f</sup>** = (rC.Abs<sup>a</sup> – rC.Abs<sup>b</sup>) – C.F.rC
**Z.Abs<sup>f</sup>** = (Z.Abs<sup>a</sup> – Z.Abs<sup>b</sup>) – C.F.Z
$\epsilon$ (M<sup>-1</sup> cm<sup>-1</sup>) Z = 57940 and rC = $1.5 \times 10^5$

**Table 3.** Calculating the number of rC and Z molecules bound to a single Gold Nanoparticle calculated using Beer -Lambert law. Molar extinction coefficient (**ε**) of PEGylated GNPs (GNP100) was used as given by the supplier (Nanopartz. Inc).

| Sample | Abs<sup>a</sup><br>SPR band<br>$\lambda = 572 \text{ nm}$ | $\epsilon$ of<br>GNP.100<br>($\text{M}^{-1} \text{ cm}^{-1}$) | Conc. of<br>GNPs<br>(M) | Cyt<br>c/GNP | Z/GNP |
| --- | --- | --- | --- | --- | --- |
| GNP100@rC | 0.972 | $1.21 \times 10^{11}$ | $0.8 \times 10^{-11}$ | 3750 | N/A |
| GNP100@Z | 1.05 | $1.21 \times 10^{11}$ | $0.86 \times 10^{-11}$ | N/A | 22906 |
| GNP100@rC@Z | 0.92 | $1.21 \times 10^{11}$ | $7.6 \times 10^{-11}$ | 1710 | 15789 |

**Table 4.** Determination of % coverage of GNP.100 by rC and Z in GNP100@rC@Z

| Sample | r<br>(nm) | S.A.rC<br>or<br>S.A.Z<br>($\text{nm}^2$) | S.A.<br>GNP<br>($\text{nm}^2$) | rC.M.<br>C<sup>t</sup> | Z.M.<br>C<sup>t</sup> | A.rC/<br>GNP<br>($\text{nm}^2$) | A.Z/<br>GNP<br>($\text{nm}^2$) | Total %<br>coverage |
| --- | --- | --- | --- | --- | --- | --- | --- | --- |
| GNP100@rC | 52 | 7.07 | 33979 | 4806 | N/A | 26512 | N/A | 78 |
| GNP100@Z | 52 | 1.53 | 33979 | N/A | 22089 | N/A | 34359 | 103 |
| GNP100@rC<br>@Z | 52 | 8.6 | 33979 | - | - | 12089 | 23683 | 104 |

**Table 5.** The calculated value of rate transfer coefficient (α) and heterogenous transfer rate coefficient (k°) using Nicholson and Shain method adapted by Lavagnini et al.^2^

| Sample | $\alpha$ | $k^\circ$ (cm.s<sup>-1</sup>) |
| --- | --- | --- |
| GNP100@rC | 0.85 | $9.5 \times 10^{-3}$ |
| GNP100@Z | 0.58 | $1.2 \times 10^{-2}$ |
| GNP100@rC@Z | 0.69 | $3.7 \times 10^{-3}$ |

**Table 6.**
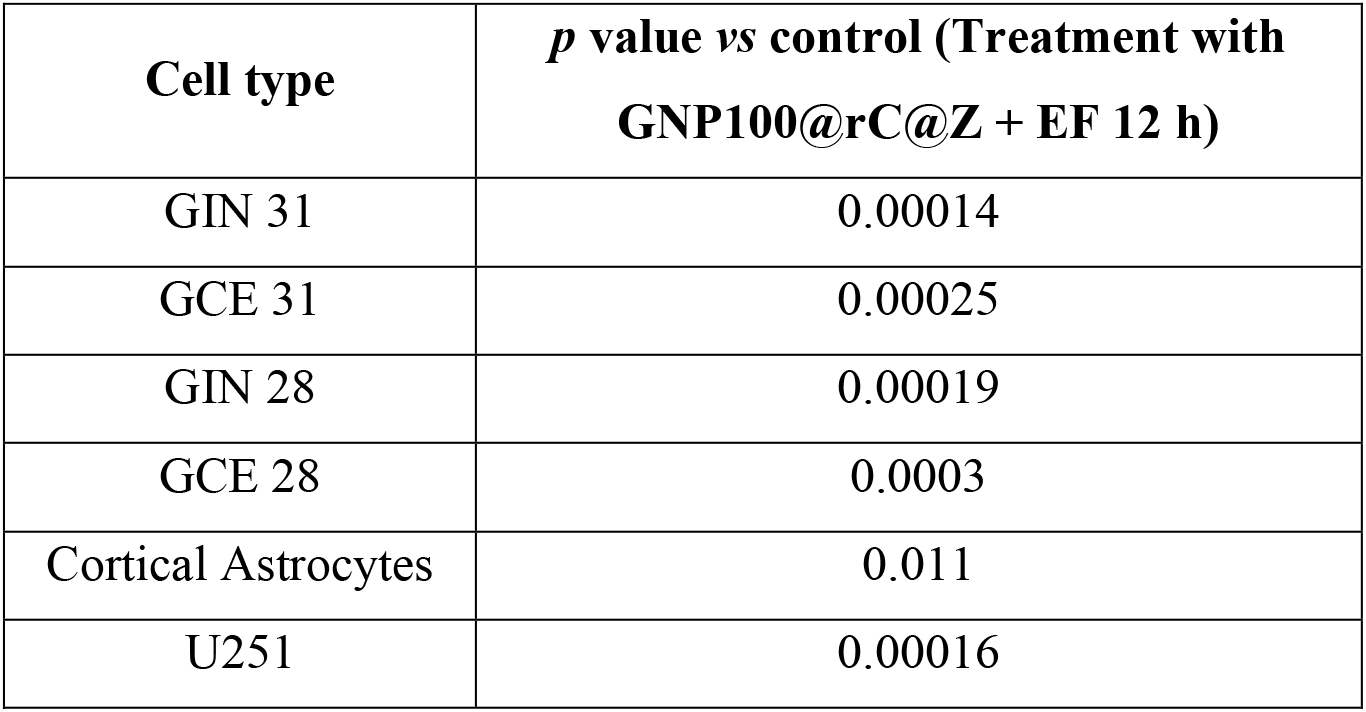
*P* values obtained from the statistical analysis of graphs shown Fig. 3 a-c (main text).

| Cell type | *p* value vs control (Treatment with GNP100@rC@Z + EF 12 h) |
| --- | --- |
| GIN 31 | 0.00014 |
| GCE 31 | 0.00025 |
| GIN 28 | 0.00019 |
| GCE 28 | 0.0003 |
| Cortical Astrocytes | 0.011 |
| U251 | 0.00016 |

